# A single-cell transcriptomic atlas of paper wasp brains highlights Kenyon cells’ role in social behavior

**DOI:** 10.64898/2026.11.05.760234

**Authors:** Colby Behrens, Martik Chatterjee, Christopher M. Jernigan, Robert D. Reed, Amy L. Toth, Michael J. Sheehan

**Affiliations:** Department of Neurobiology and Behavior, Cornell University; Ithaca, NY, United States; Department of Ecology and Evolutionary Biology, Cornell University, Ithaca, NY, United States; Department of Ecology, Evolution, and Organismal Biology, Iowa State University, Ames, IA, United States

## Abstract

Social insects provide unique opportunities for studying social interactions, but the cellular and molecular mechanisms underlying these behaviors are poorly understood. Here, we performed single-cell sequencing to profile the transcriptomes of 96,310 cells across four phenotypes of *Polistes fuscatus* paper wasps: foundresses, workers, gynes, and males, to determine how social states and social interactions are encoded in the physiology of the central brain. We characterized the cellular diversity of the *Polistes* brain and demonstrate that a substantial portion of the brain is composed of diverse populations of Kenyon cell neurons. The composition and gene-expression profiles of these Kenyon cells, as well other neural and non-neural cell populations, vary across sex, caste, and season. We also demonstrate that social interactions, and even the mere observation of a social interaction, induce significant transcriptomic responses, including signatures of neural activation, across populations of Kenyon cell neurons. Notably, the transcriptional and activational signatures elicited by observing a social interaction frequently overlapped with those induced by direct social interaction, suggesting that a shared neural program underlies both direct social interactions and social observation.

## Introduction

Social insects are powerful systems to examine the neural and physiological basis of social behavior. They exhibit division of labor among castes, which have distinct behavioral and physiological signatures despite sharing the same underlying genome^1^. The resulting social interactions can influence internal physiological states and can induce coordinated transcriptional changes in the brain across diverse neural and gene regulatory networks^2–5^. However, we currently lack a detailed molecular understanding of how social information, including social state and social interactions, are represented in brains of organisms with complex social interactions. Recent studies utilizing single-cell transcriptomics have addressed this gap, and the resulting cell atlases have provided insights into the cellular architecture, physiology, and evolution of neural tissue across vertebrates^6–9^ and invertebrates^10–12^ like social ants^13–15^ and bees^16,17^.

Primitively eusocial wasps are particularly informative models for the neural mechanisms of social behavior because reproductive status and behavioral roles are flexible throughout adulthood. In the primitively eusocial *Polistes fuscatus* paper wasp, individuals engage in complex social interactions across behaviorally distinct colony life stages^18,19^. Foundresses emerge from overwintering in the spring to initiate nests and rear offspring (Fig. 1a). After the emergence of the non-reproductive worker caste, the colony produces a final generation of reproductive individuals (gynes and males). Like other eusocial insects ^1,20,21^, *Polistes* phenotypes exhibit distinct behaviors and transcriptomic signatures^22–24^. However, caste-associated behaviors are developmentally plastic in *Polistes*, and reproductively subordinate foundresses and workers can ascend to reproductive roles following the loss of a dominant individual^25,26^. Given this flexibility, the degree of pre-imaginal biasing of caste-related physiology in *Polistes* is debated^27–29^ but likely limited compared to other highly social species like honey bees. How this plasticity is reflected in the cellular composition and physiology of the brain remains unclear.

**Figure 1.**
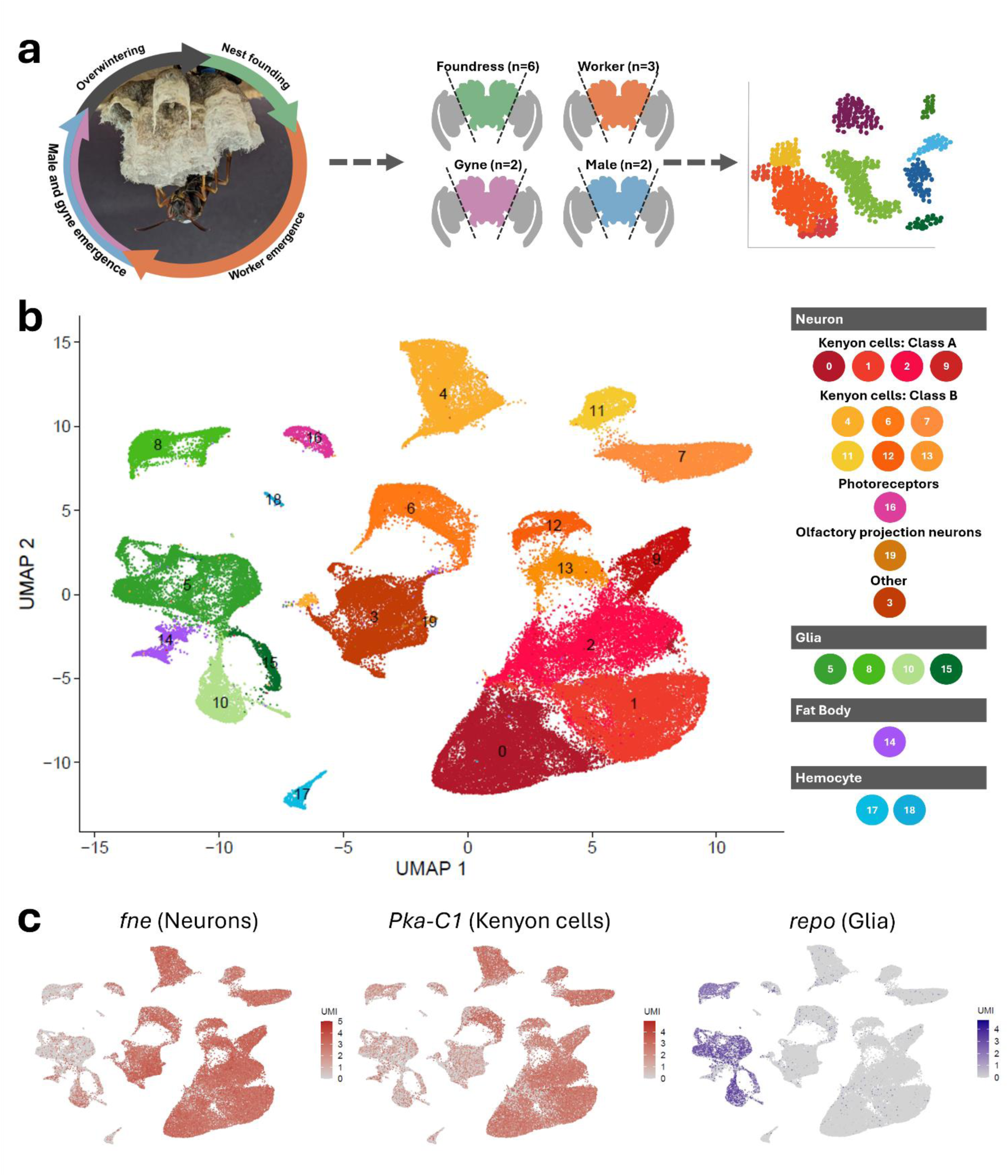
A single-cell atlas of the *Polistes fuscatus* brain. **a.** Life cycle of *Polistes fuscatus* wasps and experimental design. Foundresses build nests and care for larvae and pupae. Female workers emerge from the nest and contribute to foraging, offspring rearing, and colony growth. Later in the season, female gynes and male drones emerge. Gynes and males mate, and gynes prepare for a quiescent phase of overwintering before starting the next cycle as foundresses. Brains were collected from the four stages (foundress, worker, gyne, male) and optic lobes were removed from the central brain prior to single-cell sequencing. **b.** UMAP plot of the 20 primary clusters comprising the cell atlas. Each point represents a single cell of the 96,310 cell atlas. The legend indicates major groupings and known cell types of clusters. **c.** Marker gene expression visualized in UMAP space. The gene *fne* marks neural cells, *Pka-C1* marks Kenyon cells, and *repo* marks glial cells.

Paper wasps also exhibit complex social patterns and cognition^18,30^, including linear dominance hierarchies^31–33^, individual facial recognition^34,35^, and social eavesdropping^36^. Social interactions can trigger significant physiological changes in the central brain^37^, making *Polistes* an excellent model for dissecting how social state and experience shape brain function and composition. Though recent studies have applied single-cell transcriptomics to examine social experience in vertebrates^38,39^, comparable cellular-resolution investigations are largely unexplored in invertebrates.

Here, we generated a comprehensive single-cell transcriptomic atlas of the *Polistes fuscatus* brain. We used this resource to identify social phenotype-related patterns of neural composition and gene expression. We then examined how direct and observed social experience influences transcriptional states across neural cells.

## Results

### Cell classification of *P. fuscatus* central brains

To generate a cellular-resolution transcriptomic atlas, we performed single cell transcriptomics on replicates of central brains harvested from foundress (n=6 libraries), worker (n=3), gyne (n=2), and male (n=2) wasps (Fig. 1a; 5-6 wasp central brains/library). After filtering and integration, the atlas retained 96,310 cells and 11,324 genes. Filtered cells were clustered at two levels, primary (resolution = 0.2) and secondary (resolution = 0.8), to enable analysis of major cell types and their subpopulations. This resulted in 20 primary clusters (Fig. 1b) and 39 secondary clusters (Fig. S1a). The following analyses focus on the central brain to the exclusion of the optic lobe to focus on higher-order integration centers. However, we sequenced worker optic lobes to provide a whole brain atlas for wasps. The combined worker central brain-optic lobe atlas is presented in the supplemental material (Fig. S2, Table S3, S6).

Clusters were annotated with canonical insect marker genes^10,13,14,16^ and cleanly separated major cell types (Fig. 1c; Fig. S1b; Table S4). Thirteen clusters were annotated as neurons according to co-expression of neural markers (*fne, brp*, *nSyb*, *Syt1*), including 10 Kenyon cell clusters (*Pka-C1, CaMKII, trio*), an olfactory projection neuron cluster (OPN; *acjc, ct)*, and a photoreceptor cluster (*ninaC, ninaB*), which likely originated from ocelli retina that were not removed during brain dissection. The 10 Kenyon cell clusters are especially notable as they collectively represent 70.61% of the atlas, substantially more than central brain atlases of *Harpegnathos* ants (36%) and *Drosophila* flies (10%)^13^. In insects, higher order cognitive processes like memory formation and learning occur primarily in mushroom bodies and their Kenyon cell neurons^40–42^, which drive these cognitive functions, regulate social behaviors, and facilitate social memory^43,44^. Morphological measurements suggest that mushroom bodies are large in wasps^45^ and these single-cell data reinforce the idea that mushroom bodies, and potentially cognitive abilities, are expanded in this species compared to other insects^13^.

Four glial clusters (*repo*, *bdl*) were also recovered, including astrocytes (*Gs2, Gat)* and cortex glia (*repo, bdl, wrapper*), and two hemocyte clusters were identified (*Hml, Fer2LCH*; Fig. 1b). We also identified a cluster of head fat body cells (*ACC, svp*), which have been previously implicated in metabolism, immune response, and courtship behavior in *Drosophila*^46–48^. Head fat tissue abundance can vary across insect species and is rarely identified in single cell datasets (but see Croset et al. 2018). The clear presence of a fat cluster suggests that *Polistes* wasp heads may be fattier than other insects, consistent with our experience dissecting wasp heads compared to those of other social insects (CMJ, personal observation).

Neurotransmitter markers further characterized neural cells. Cholinergic neurons (*VACht*) were distributed across multiple neural clusters (0, 4, 6, 7, 11, 12, 19) and glutamatergic neurons (*VGlut*) were localized to clusters 3 and 19. Markers for GABAergic (*Gad1*), monoaminergic (*Vmat*), and dopaminergic neurons (*DAT, ple*) were expressed in neuronal cluster 3 and fat body cluster 14 (Fig. S3a-c). Head fat bodies exhibit similar patterns in *Drosophila*, where they express *Vmat*^49^ and utilize the dopaminergic pathway to convey immune response signals to mushroom bodies^50^. Indeed, *Polistes* Kenyon cells were enriched for dopaminergic and other neurotransmitter receptors (Fig. 3d), indicating that the interactions between Kenyon cells and peripheral tissues like the fat body is a fruitful topic for future investigation.

To evaluate the robustness of the dataset, we mapped *Polistes* clusters against single-cell datasets from three other social insect species (*Monomorium pharaonis*^13^, *Harpegnathos saltator*^14^, *Apis mellifera*^16^) and *Drosophila melanogaster*^10^ and assessed transcriptional similarity via MetaNeighbor AUROC scores^51^. To reflect biological structure, conserved cells should cluster by cell type, not species. Indeed, Kenyon cells, glia, and optic lobe neurons consistently clustered according to cell type across each species comparison (Fig. 2a, S4-S6). Additionally, MetaNeighbor analyses recapitulated the previously described class A and class B Kenyon cell subgrouping across all examined species^13^ (Fig. 2b). Transcriptomic-defined class A cells are associated with memory formation and learning, while class B cells are associated with locomotion and circadian regulation^13^. Wasp class A Kenyon cells preferentially expressed *EipS3f*, a marker characterizing class I large type Kenyon cells in bees, while class B cells preferentially expressed markers *Rbpc* and *Cow*. Class B clusters 4 and 7 also preferentially expressed the neuropeptide *sNPF*. Marker genes were also enriched for GO biological processes matching known functions in the central and whole brain atlases (Fig. S7). For example, neurons were enriched for synaptic signaling and glia were enriched for lipid and carbohydrate metabolic processes.

**Figure 2.**
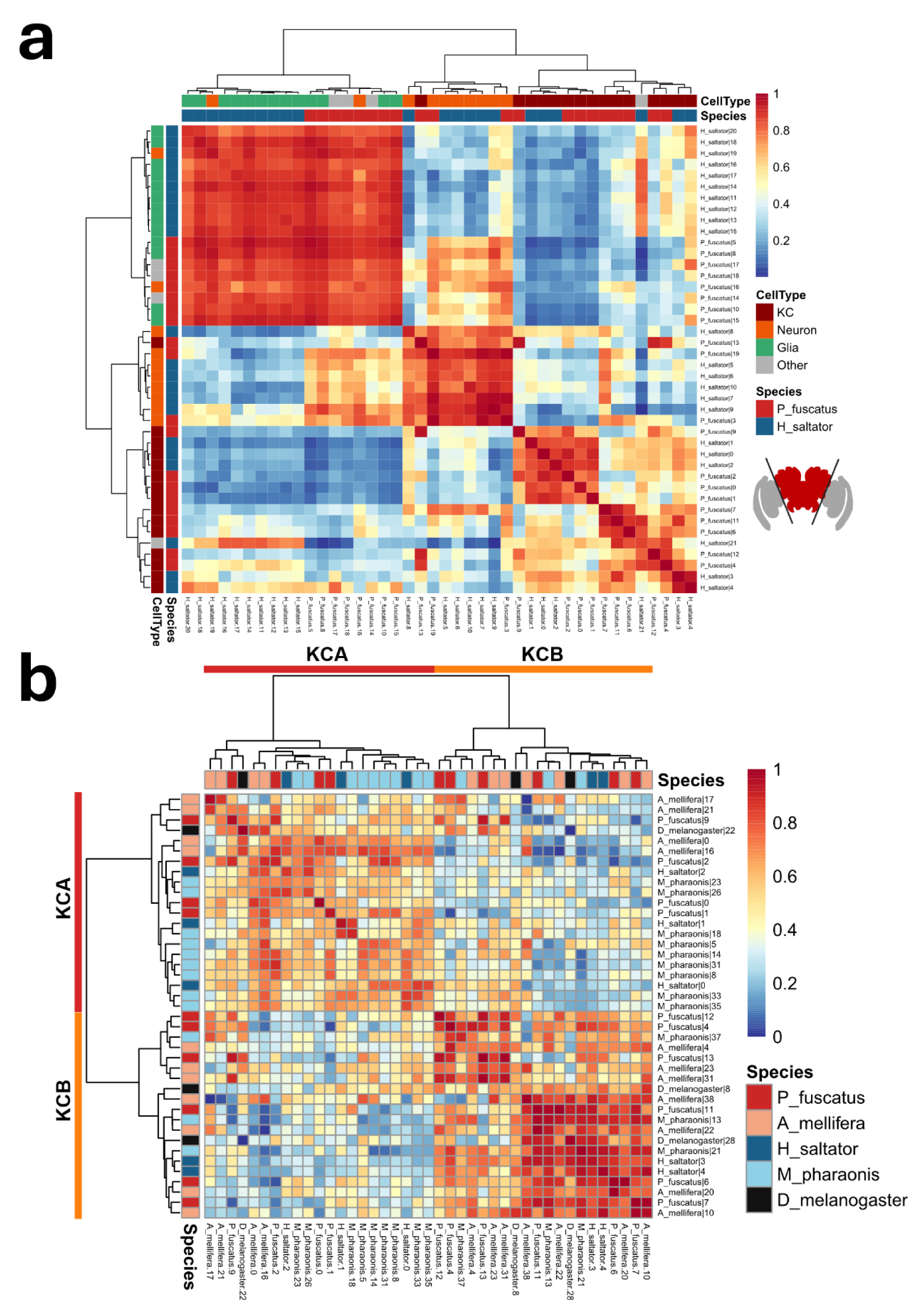
The *Polistes* brain atlas is robust and recapitulates known Kenyon cell subtypes. **a.** Pairwise transcriptional similarity of central brain cell clusters from *Polistes fuscatus* against the jumping ant *Harpegnathos saltator* from MetaNeighbor AUROC scores. Major cell types generally clustered together across species, though *P. fuscatus* cluster 13 was notably separated from other Kenyon cells clusters. Columns and rows are annotated according to the species and cell type. KC, Kenyon cell. **b.** Class A and class B Kenyon cells cleanly separate based on transcriptional similarity (AUROC scores) of Kenyon cell clusters from *Polistes, Drosophila, Harpegnathos, Monomorium,* and *Apis* datasets. The heatmap color has been rescaled to enhance contrast and better reflect dendrogram relationships.

### Cellular compositional differences across phenotypes

Primary and secondary cell clusters were shared across all adult phenotypes (Fig. 3a). 50% of primary clusters (10/20) exhibited differential cell count proportions between at least two phenotypes (Fig. 3b; Tables S7-12), and 69% (27/39) of clusters exhibited differential cellular abundance when analyzed at the more granular secondary clustering level (Tables S13-18). Neural investment correlates with behavioral and sensory specialization,^52,53^ and relative shifts in cluster abundance across groups (Fig. 3) indicate phenotype-associated differences in brain cellular composition, as discussed below.

**Figure 3.**
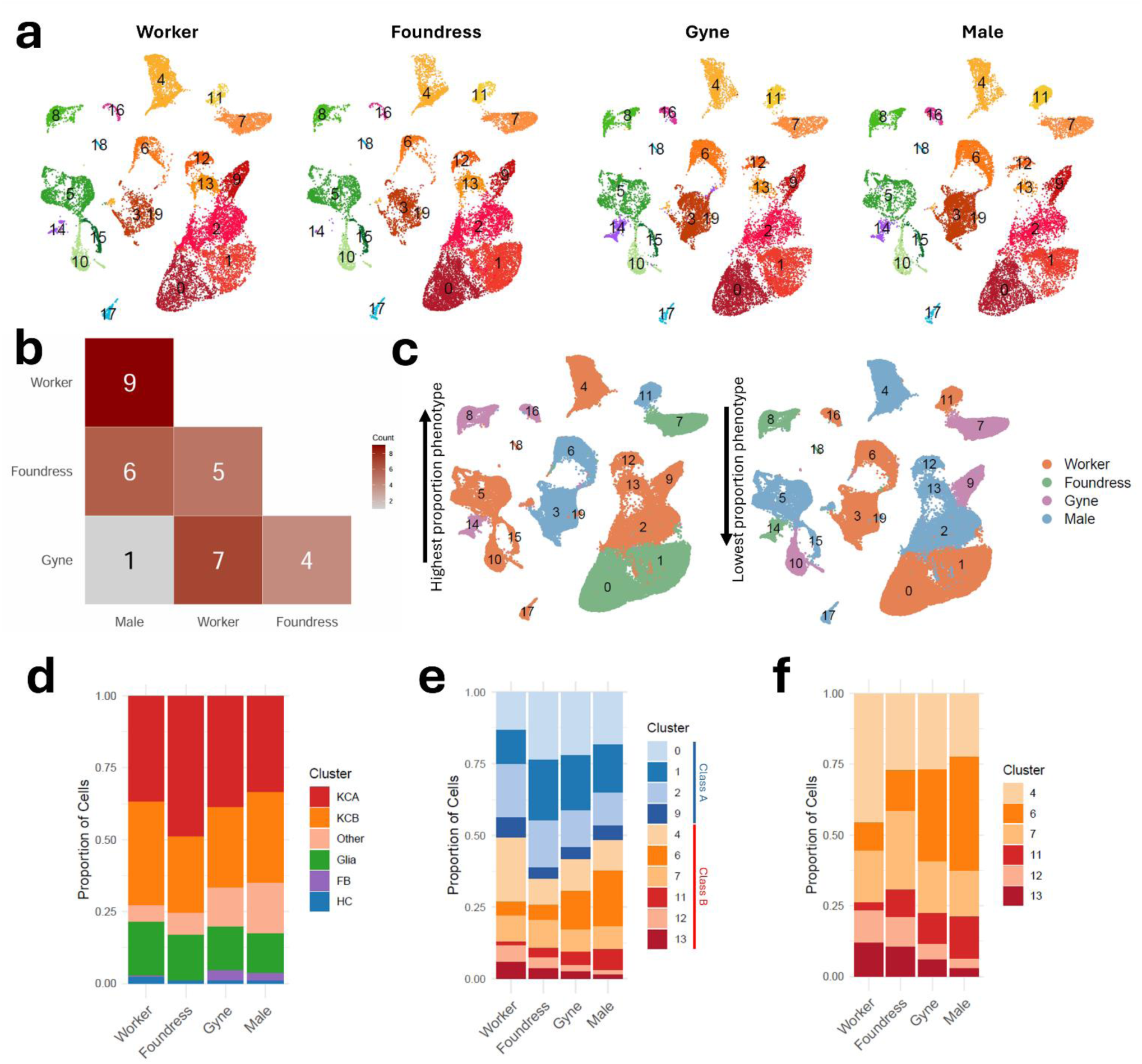
Wasp phenotypes exhibit distinct brain composition. **a.** Though cell types are shared, differences in cellular abundance of multiple clusters (e.g. 6, 14) are readily apparent. Cells from gynes, foundresses, workers, and males are plotted in UMAP space. Cells from each stage have been down sampled to visualize the relative abundance of major cell types. **b.** The number of clusters with significant differential cell abundance between phenotypes, as measured by scCODA. Gynes and males exhibited the fewest compositional differences. **c.** UMAP representations of cell clusters colored according to the phenotype with the greatest proportional representation or the least. Workers exhibit the extreme proportions (highest or lowest) in 17/20 clusters. **d.** Wasp foundresses have proportionally higher Kenyon cell counts than other wasp phenotypes. The stacked bar chart displays the abundance of major cell types against the total number of cells in the brain for each phenotype. KCA, Kenyon cell class A; KCB, Kenyon cell class B; FB, fat body; HC, hemocyte. **e.** Within Kenyon cells, wasp phenotypes differ in the abundance of cell clusters. For example, KC cluster 4 is expanded in worker brains compared to other phenotypes. The stacked bar chart displays the proportional representation of Kenyon cell clusters out of the total Kenyon cells in each phenotype. **f.** Within class B Kenyon cells, cluster proportions differ across phenotypes, suggesting that these patterns are not driven by the abundance of KCA cells or neurons overall.

Each wasp phenotype exhibited the highest or lowest relative abundance for at least one primary cell cluster (Fig. 3c). Workers differed the most from other groups with 21 of 60 pairwise comparisons (based on proportional cluster abundances) showing significant differences. In contrast, gynes differed the least from others, with only 12 pairwise cluster abundance differences, while foundresses and males each differed in 16 pairwise comparisons. The relatively small number of differences involving gynes reflects their high similarity to males, as the relative abundance of only one primary cluster differed between males and gynes, whereas nine primary clusters differed between males and workers. Sexual brain dimorphism is expected to underlie sex-specific behaviors in other insects^13,54^, so the apparent minor compositional differences between similarly-aged reproductives (gyne and male) in paper wasps is striking. These similarities may be partially driven by seasonal effects. Gynes and males exhibited greater relative representation of fat body cell clusters compared to workers and foundresses, mirroring patterns in abdominal fat content^55,56^. Behavioral differences among phenotypes may also drive aspects of brain compositional differences. Males and gynes had the highest relative proportion of ocellar photoreceptors, and this difference may reflect the fact they both disperse from colonies in a seasonally dependent manner that may rely on their ability to detect differences in day length. Males, followed by gynes, possessed more abundant cluster 3 cells, which contains monoaminergic neurons, as well as two different clusters of Class B Kenyon cells (6 and 11). How these neurons may contribute to male specific behaviors or mating more generally is unknown. Further evidence for neural specialization is seen in foundresses and workers, as foundresses exhibited the highest proportion of large Class A Kenyon cells (clusters 0 and 1) and a Class B Kenyon cell cluster (7). All other clusters had the greatest proportional representation in workers (Fig. S8).

Wasp phenotypes exhibited distinct Kenyon cell proportions relative to other cell types, as well as shifts in Kenyon cell composition, as expected from anatomical studies^57–59^. Foundresses and workers exhibited the highest proportion of Kenyon cells in the brain, suggesting that cognitive demands during foraging, brood care, and social interactions may be higher than in gynes and males (Fig. 3d). Class A Kenyon cells are proportionally expanded in foundresses compared to other groups, potentially facilitating enhanced memory consolidation and sensory integration. Class A clusters 0 and 1 were relatively less represented, whereas class B cluster 4 was relatively more represented in worker brains, but the functions of these clusters are currently unclear (Fig. 3e-f). Though gynes develop into foundresses after overwintering quiescence, they exhibited substantial divergence in the relative abundance of Kenyon cell classes. For example, class A Kenyon cells are less abundant in gynes. This pattern appears to be driven by cluster 6, a Class B Kenyon cell subpopulation that is notably expanded in gynes and males. If changes in gyne and foundress neural composition are driven primarily by cell death, rather than neurogenesis, then the decrease in cluster 6 could indicate preferential pruning associated with the shift from fall mating to spring colony foundation^52^.

### Gene expression differences across phenotypes

Wasp brains exhibited a wide range of phenotype-specific gene expression differences (Fig. 4a, Table S19) that varied across clusters (Fig. 4b). Differentially expressed gene counts scaled with cluster size due to increased statistical power (Fig. S9), so we examined overall patterns across secondary clusters, which provided more balanced cell numbers across clusters. A substantial portion of the phenotypic divergence reflected sex-biased transcription, as thousands of genes were differentially-expressed across sexes (Fig. 4a). For example, the transcription factor *fruitless*, which regulates sex-specific behaviors in insects^60,61^, was consistently female-biased across *Polistes* neural clusters (Fig. 4c). *Dop2R*, a dopamine receptor, was similarly female-biased across neural clusters. Dopamine signaling is associated with female dominance and caste-specific behavior in *Polistes* and other insects^55,62^, and this suggests that male neural populations may be less sensitive to dopaminergic cues.

**Figure 4.**
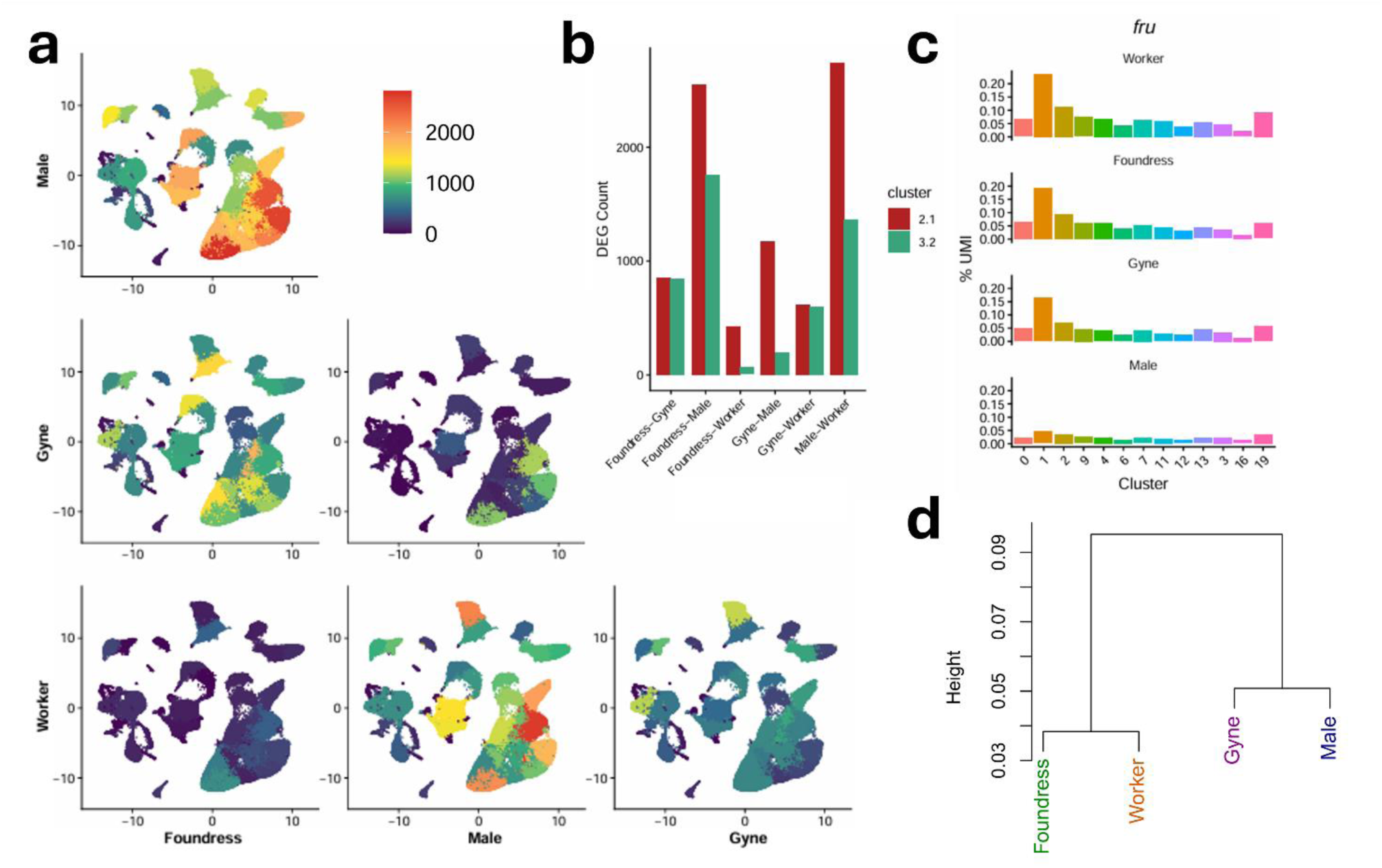
Wasp phenotypes exhibit distinct transcriptional profiles across clusters. **a.** Counts of cluster-specific differentially expressed genes (adjusted p value < 0.05, log2FC > 0.5) are plotted across each pairwise contrast. Contrasts against males nominated the greatest number of DEGS, particularly in Kenyon cell clusters. Cell clusters are plotted in UMAP space. **b.** Kenyon cell subcluster 2.1 exhibits greater differential expression than the similarly-sized neural cluster 3.2, suggesting that variation in DEG numbers is not due to cluster size alone. **c.** Expression levels (% of cluster UMIs) of the transcription factor *fruitless* across neural clusters. Female phenotypes exhibit higher expression than males across clusters. **d.** Brain-wide hierarchical clustering of the 3,000 most variable genes indicate that early season stages (foundress, worker) are transcriptionally distinct from late season stages (gyne, male).

Caste and season also contributed to transcriptional divergence. *Foraging* (*for*), a gene encoding a protein kinase associated with worker-like phenotypes in other social Hymenoptera^63^, was consistently upregulated in worker cells. We also detected phenotype-specific patterns of neurotransmitter expression. For example in neural cluster 3, foundresses and workers expressed higher levels of *VGlut*, a marker of glutamatergic excitation, while gynes and males expressed more of the inhibitory *Gad1* gene. This suggests that the phenotypes engaged in foraging and brood care exhibit more excitatory neural profiles, while stages that primarily engage in reproduction exhibit inhibitory, GABAergic-driven profiles. The change in glutamatergic/GABAergic ratios between the gyne and foundress stages further highlights neurodevelopmental changes, but excitatory/inhibitory expression is known to vary with age^64^, so these patterns likely reflect a mix of season, behavior, and age effects.

Finally, pseudobulk analysis suggests that behaviorally-similar groups (e.g. brood-provisioning foundresses and workers) share more similar transcriptomic profiles than behaviorally distinct phenotypes (e.g. non-provisioning gynes; Fig. 4d). This is consistent with results from previous molecular studies,^22^ but these patterns are heterogeneous across clusters (Fig. S10) and suggest that subtle patterns are likely obscured under bulk sampling.

### Merely observing a social interaction elicits socially-mediated gene expression in Kenyon cells

Paper wasps make extensive use of visual signaling to mediate social interactions^19^. In *P. fuscatus*, females visually recognize other wasps^34^, remember their past interactions^35,65^, and are capable of assessing conspecifics via social observation alone^36^. We previously hypothesized that learning through merely the observation of third-party social interactions is likely to co-opt the neural mechanisms used to process first-party social interaction in wasps, with Kenyon cells as likely candidates^66^.

To examine the transcriptional responses of brain cells when wasps engage in or observe a social interaction, we profiled single cells from foundresses across three social conditions (Fig 5a). Paired ‘fighters’ engaged in a direct first-party social interaction. ‘Watchers’ were placed alone in an adjacent area with visual access to the interacting pair. A third group of solitary control wasps were isolated in similar arenas to the watchers but without visual access to other wasps. Differential expression between fighters and control wasps reflected transcriptomics shifts associated with first-party interaction, similar to previous studies^37^, while differences between watchers and controls reflected gene expression associated with observation of third-party social interactions. Shared expression patterns in fighters and watchers, relative to controls, indicated cluster-specific responses common to direct and indirect social interactions.

**Figure 5.**
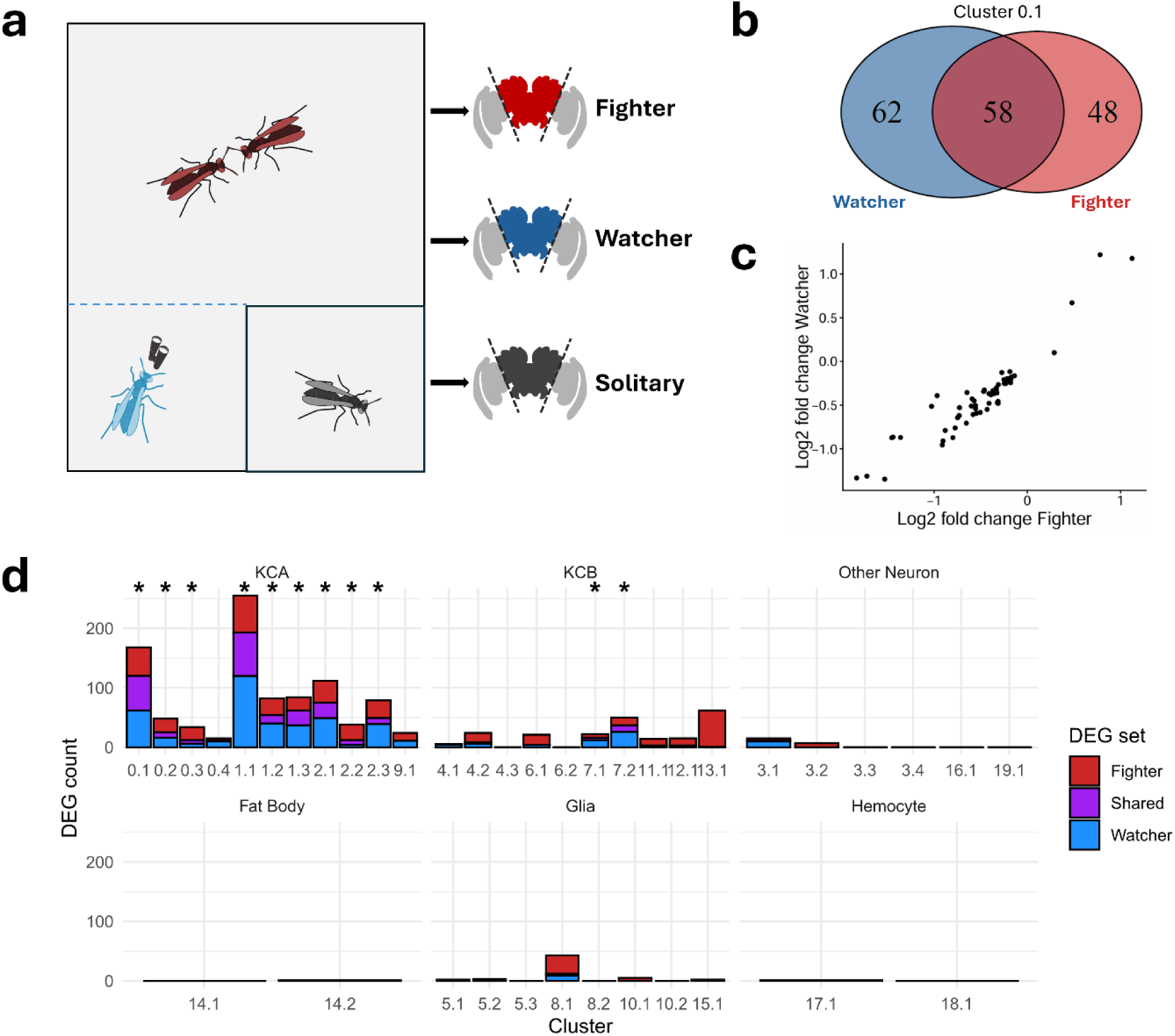
Observation of a social interaction is sufficient to elicit socially-mediated gene expression patterns. **a.** Experimental design of the social experience assay. Foundresses were assigned to one of three social conditions. “Fighters” engaged in first-party physical interactions in a small arena, “watchers” experienced a third-party observation of a physical interaction through a clear barrier, and “solitary” wasps were placed individually in arenas without any social stimuli. Afterwards, each wasp’s central brain was dissected and processed for single-cell RNAseq. **b.** Fighters and watchers exhibit significant overlap in differentially expressed genes in cluster 0.1 (hypergeometric test; 58 genes, p < 0.01). **c.** Shared DEGs from cluster 0.1 exhibit concordant expression compared to the solitary condition, suggesting that the transcriptional response to an observed or experienced social stimulus is similar. **d.** Across clusters, foundresses exhibited significant differential expression (adjusted p value < 0.1) in response to engaging in or viewing a social interaction, with the majority of DEGs detected in class A Kenyon cells. Stacked bar plots display the number of DEGs for each cluster, faceted by cell type. Asterisks indicate significant DEG overlap across fighter and watcher conditions.

Multiple distinct lines of evidence suggest that third-party social observation and first-party social interaction activated shared transcriptional responses, especially in Kenyon cells. First, we estimated cluster-wise differential expression between each social treatment and the solitary control condition. Class A Kenyon cells exhibited the greatest transcriptional responses to social treatments, matching prior work demonstrating that mushroom bodies respond to social stimuli^5,67^. Notably, fighters and watchers exhibited a high degree of overlap and concordance across several clusters (Fig. 5b-c). Hypergeometric overlap tests identified 11 clusters where fighter and watcher differentially expressed gene (DEG) sets significantly overlapped (Fig. 5d). Notably, genes differentially expressed in both social conditions tended to shift in the similar direction relative to solitary wasps, suggesting that viewing a third-party social interaction can elicit a similar response to direct first-party interactions. Intriguingly, surface glial cluster 8.1, analogous to the blood brain barrier, also exhibited significant differential expression in fighters, but not watchers. Surface glia have been previously implicated in regulating social states in ants^15^, and this suggests that direct interactions with other wasps may alter how the brain interacts with hemolymph in paper wasps.

Second, to link transcriptional changes to neural activity, we examined cluster-specific activity using immediate early gene (IEG) expression. Neural excitation induces transcriptional waves across cells, and initial waves are characterized by expression of IEGs^68,69^. IEG expression is often sparsely detected in single-cell datasets, but panels of co-expressed IEG-like genes can be used to estimate patterns of neural excitation^8,70^. Two canonical insect IEGs^69,71^ (*hr38*, *sr*), were strongly co-expressed across wasp neurons (Fig. 6a), while two other IEGs (*kayak*, *Jra*) were not (Fig. S11a). We therefore used cluster-wise co-expression with *hr38* and *sr* to identify 18 highly co-expressed genes with putative IEG-like expression patterns (Fig. 6b). This panel indicated that neural clusters exhibit substantially higher activation than non-neural clusters, suggesting that the panel captured neural excitation (Fig. 6c).Having established an activation signature, we next asked which genes are preferentially expressed in putatively-activated vs non-activated cells (see Methods) to identify gene-defined subpopulations that respond to social stimuli^70^. For each social condition, we identified DEGs between activated and non-activated cells, which enabled identification of neural subpopulations whose activation was specific to a first-party interaction, third-party observation, or shared by both.

**Figure 6.**
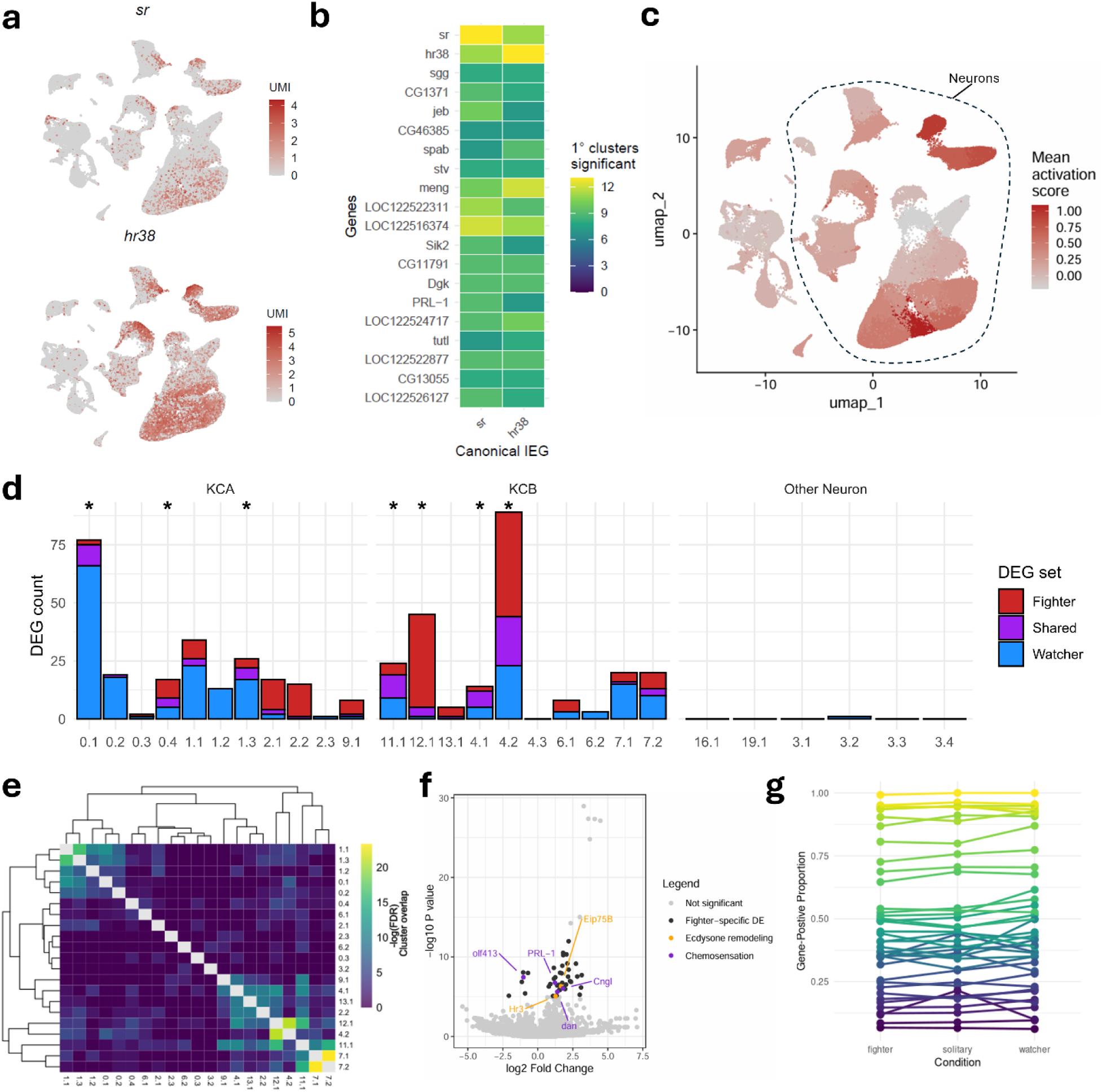
Social interactions are encoded in the activational state of Kenyon cell neurons. **a.** Expression of canonical IEGs *sr* and *hr38* plotted in UMAP space. Expression is primarily limited to neural clusters. **b.** A panel of eighteen IEG-like genes were co-expressed with canonical IEGs *sr* and *hr38*. IEG-like genes were classified by upregulation in IEG-positive cells compared to IEG-negative cells in the majority of neural clusters. Genes without known *Drosophila* orthologs are named with *Polistes* LOC identifiers. **c.** The mean activation score of secondary clusters, as assessed by the panel of IEG-like genes, plotted in UMAP space and demonstrating that neural clusters exhibit higher average activation than non-neural clusters **d.** Activation-associated DEGs were almost exclusively found in Kenyon cell neurons. The social conditions exhibited greater overlap of DEGs than expected by chance across 7 clusters, suggesting that the similar neural populations are activated in respone to social stimuli. The number of activation-associated DEGs from each condition are plotted. No activation-associated DEGs were identified in non-neural clusters. Asterisks indicate significant overlap across fighter and watcher conditions. **e.** Shared first-party/third-party DEG sets from each cluster were compared pairwise using hypergeometric overlap tests. Heatmap values indicate the significnance of overlap between DEG sets. Activation-associated DEGs were primarily non-overlapping across clusters, suggesting that transcriptionally distinct populations of cells are activated in each neural cluster. **f.** Activated neurons in cluster 12.1, which responded strongly to first-party physical interactions, are enriched for genes related to chemosensation and ecdysone remodeling. The volcano plot displays the log2 fold change of genes between activated and non-activated cells of cluster 12.1. **g.** The proportion of cells expressing cluster 12.1 DEGs was consistent across conditions, suggesting that the observed patterns are the result of differential activation, not differences in cell composition. Each line plots the number of cells in the cluster that expresses a DEG.

Hundreds of genes were preferentially expressed in activated cell populations in response to social stimuli. The majority of this signal originated from Kenyon cell clusters (Fig. 6d), further underscoring the mushroom bodies’ role in regulating social behavior^44,67^. Like the overall transcriptional response, both social conditions exhibited substantial overlap in activation patterns. Hypergeometric tests suggest that 7 clusters exhibited significant overlap in gene-defined cell population activation in response to direct physical interaction and during social observation, suggesting that these populations responded to both forms of social experience. Although activational signatures were more similar than expected by chance between watchers and fighters within multiple clusters, activation-related DEGs were distinct to individual clusters and tended to show little to no overlap between clusters (Fig. 6e). This means that distinct genes defined the neural populations with shared activation across clusters. This analysis shows that participating in or observing a social interaction engaged shared cell-type specific neurogenomic responses in paper wasps. First-party social interactions are known to elicit excitatory and physiological responses in honeybee Kenyon cells^5,44^, and these results implicate a role for Kenyon cells in third-party social observation as well in paper wasps.

In addition to visual signals, wasps rely heavily on olfaction for social communication and utilize non-volatile cuticular hydrocarbons (CHCs) to mediate recognition and social behaviors^72,73^. We predicted that cell populations that specifically activate following first-party interactions, but not visual-only observations, would be enriched for chemosensory processes. Consistent with our prediction, DEGs from cluster 12.1 activated neurons (Fig. 6f), which responded strongly only in the fighter condition, were linked with chemosensation, including *olf413*, *dan*, *Cngl*, and *PRL-1*^74–77^. Cells expressing canonical ecdysone-related transcription factors (e.g. *Eip75B*, *Hr3*) were also activated alongside genes associated with synaptic function (e.g. *cac*, *KaiR1D*), suggesting a link between first-party interactions and ecdysone-mediated olfactory remodeling. These gene-defined cell populations were preferentially activated in the fighter group but were expressed in similar proportions across all three conditions (Fig. 6g), consistent with differential activation of neural populations rather than changes in cell abundance.

## Discussion

### Cellular architecture of the Polistes brain varies across phenotypes

This study utilized scRNA-seq to generate a single-cell atlas of the *Polistes fuscatus* brain and characterize neural and non-neural cell populations across a range of social states and conditions. The cellular composition and regulatory dynamics of the brain varied significantly across wasp castes, sexes, and life-history stages. This pattern was particularly apparent across mushroom body Kenyon cell clusters, which dominated the cell atlas compared to other insect species^10,13,15,16^. Kenyon cell abundance has been associated with greater cognitive ability and behavioral innovation across insects^78,79^. We found substantial phenotype-associated variation across Kenyon cell subpopulations, suggesting that differences in cognition across wasp phenotypes may be reflected in mushroom body composition. Indeed, although workers and foundresses are both capable of cognitive tasks like individual facial recognition^80^, workers exhibit poorer performance and shorter memories than foundresses^81^. The expansion of Kenyon cells in this atlas is consistent with morphological studies^59^, and further investigation linking Kenyon cell subpopulations to cognitive function in *Polistes* is warranted.

Our atlas also revealed phenotype-associated signals in non-neuronal cell populations. Glia, known for developmental and metabolic functions^82^, are increasingly recognized for contributing to behavioral processes like the sleep-wake cycle^83^ and caste dynamics^14,15,84^. Similarly, peripheral tissues like head fat bodies modulate pathogen avoidance in *Drosophila*^50^. Here, wasp phenotypes were also differentiated by the composition and expression of non-neural clusters. Physiological profiles vary across social insect castes^21,22^, and our results suggest that these differences are driven by heterogenous patterns across the cellular landscape of the brain.

### Kenyon cells are implicated in regulating and interpreting social behavior

Social experiences are known to generate long- and short-term effects on physiology and behavior, but how social experiences are encoded in insect brains is still poorly understood. We compared expression and neural activation in samples that physically engaged in direct social interactions versus those that only observed third-party interactions. Social exposure induced significant differential expression in the brain, as has been observed in other systems^3,5^, but this response was almost entirely localized to Kenyon cell clusters. Additionally, IEG analyses indicated that specific subpopulations of Kenyon cells were activated following social stimulation, suggesting that neural responses to social experiences are distributed across distinct neural subpopulations throughout the mushroom body, rather than being confined to a single cell cluster. Mushroom bodies have been previously implicated in encoding memories related to social interactions^5,44,67^ and a myriad of other behavioral paradigms^43,85,86^. Our results recapitulated this pattern but also indicate that Kenyon cells neurons experience consistent activation and physiological shifts after merely observing a social interaction. Kenyon cell subtypes that responded to first-party interaction were frequently engaged in the response to third-party observation as well. Our differential expression and differential activation analyses have distinct sets of analytical approaches and controls yet find the same overall pattern of greater sharing than expected between the wasps engaged in first-person social interactions versus third-party observation. The consistent patterns across different analysis approaches suggest our finding of shared neural activity across conditions is robust.

The data presented here show clear overlap in neurogenomic responses between wasps that engage in social interactions and those that merely view the interaction. This may simply be driven by shared neurogenomic responses to seeing another wasp. This interpretation would be consistent with behavioral and neural studies of individual face recognition in paper wasps^36^. Electrophysiological recordings in wasps suggest that cells roughly analogous to the face cells of primates^87^ are in the lateral protocerebrum^88^ rather than the mushroom bodies where Kenyon cells are located. In light of the electrophysiological data, the shared patterns of Kenyon cell expression and activation observed here implicate Kenyon cells as a component of the neural circuitry possibly involved in integrating identity information with social valence^66^. We cannot rule out the possibility, however, that these shared responses reflect the act of engaging in an interaction and interpreting others’ interactions make use of a shared neural substrate. Recent work in mice has identified neurons that are active during aggression and specifically during observations of aggressive acts^89,90^. Similar neurons that mirror aggression (or other aspects of social interactions) in wasps would help explain the observation that this species can interpret relative dominance outcomes from a contest that they have merely observed^36^. While these data suggest the potential for neurons that mirror activity and interpretation in an insect, additional expression and electrophysiological studies will be needed to understand how wasp brains interpret social interactions.

Additionally, the observed response to direct social interactions is likely an underestimate of the true responses. Arguably, the “watcher” wasps had a relatively more consistent experience compared to “fighter” wasps, where individuals were sampled from winners and losers of their contests. Our prior work has shown that initial neurogenomic responses in fighting wasps (i.e., those sampled here) are similar regardless of dominance status, with winner and loser effects emerging hours later^37^. Winner and loser effects can produce distinct behavioral and physiological patterns across taxa^91–93^, including paper wasps^37^, and heterogeneous responses to winning and losing may obscure other patterns associated with direct interactions. Future studies of these paper wasps have the potential to examine how both social exposure and social assessment are biologically encoded in the brain and can provide further insights into the circuitry involved in insect social processing.

## Methods

### Brain dissection, cell isolation, and sequencing

Male and gyne wasps were collected in Tompkins County, New York, in 2022. Worker and foundress wasps were collected in 2023. Animals were sacrificed via rapid decapitation and dissected immediately under ice-cold 1× Sf-900 III SFM medium (Thermo Fisher Scientific). Optic lobes were separated from the central brain but peripheral connective and fat tissue were not removed. Following dissection, brains were cut into approximately 200 x 200 micron pieces and pooled according to treatment. Each brain dissection was completed in under 5 minutes, so a pool of 5-6 brains was processed for dissociation in approximately 20 minutes. Once brains were pooled, the medium was replaced with 750 µL of papain (lyophilized; Worthington Biochemical) suspended in EBSS at a final concentration of 2 mg/mL. Brains were gently triturated by pipetting up and down approximately ten times to dissociate the tissue without lysing cells. The samples were then incubated in the papain solution for 30 min at 37°C with rotation at 800 rpm in an incubator shaker. During incubation, the tissue was gently pipetted every 10 minutes to further aid dissociation.

After incubation, the cell suspension was centrifuged at 300 × g for 5 minutes. The supernatant was removed and the pellet was resuspended twice with 1 mL of 1× PBS containing 0.04% BSA (Thermo Fisher Scientific) and RNase inhibitor (MilliporeSigma), with centrifugation at 300 × g for 5 min at 4°C between suspension. After the final centrifugation, the supernatant was removed and the pellet was resuspended in 500 µL of PBS/BSA/RNase inhibitor solution. The suspension was then passed through a 20 µm pluriStrainer to remove debris. Cells were counted using a Countess 3 automated cell counter (Thermo Fisher Scientific) to assess cell number and viability. Cell viability was 90-95% across all treatments.

Single-cell libraries were prepared using the Chromium Single Cell 3′ Reagent Kits v3.1 (10x Genomics) by the Biotechnology Resource Center BRC Genomics Facility (RRID:SCR_021727) at the Cornell Institute of Biotechnology. Libraries were sequenced on an Illumina NextSeq 2000 P3 flow cell at the Cornell Genomics Facility.

### Read processing

FASTQ files were processed with CellRanger v9.0.1 pipeline, which performed all alignment, barcode processing, and UMI counting to quantify gene expression. Reads were aligned against the *P. fuscatus* genome^94^. We updated the NCBI gene models by generating a library of PacBio IsoSeq data for wasp heads and running the software PASA on the consensus reads^95^. The genome annotation files used in this work are provided as supplemental materials. After CellRanger processing, raw and filtered gene-barcode matrices files were input into SoupX^96^ to estimate and remove ambient RNA contamination. Background-corrected cell expression profiles were then used in downstream analyses. Cells with less than 250 or greater than 2,500 features were discarded to reduce the risk of doublets or low-quality cells. Cells with elevated mitochondrial expression are often considered contaminated, apoptotic, or otherwise low-quality. We therefore removed cells with >20% mitochondrial RNA, matching thresholds from other insect brain studies^97,98^. Across all libraries, we retained 96,310 central brain cells.

To characterize the transcriptomic profile of the *Polistes* whole brain, both central brains and optic lobes were sequenced from worker samples. Optic lobes were separated from the central brain and three libraries (6 individuals / library) were processed for scRNAseq as above via CellRanger and SoupX pipelines. After processing, optic lobe libraries were merged with worker central brain libraries. The central brain and optic lobe datasets were derived from distinct tissues, so we opted for merging rather than integration to avoid overcorrection of tissue-specific signal. The resulting 33,439 cells of the worker whole brain atlas were clustered at resolution = 0.8, which captured major cell populations from both tissue types. We focus our analyses on the central brain to focus on the non-visual, integration centers of the brain, but provide optic lobe data as a reference for future studies.

### Dimensional reduction and clustering

After filtering, we used Seurat v5.3.0^99^ for normalization, integration, clustering, and visualization. Libraries were normalized with SCTransform. To account for potential batch effects, normalized values from all samples were integrated into a single atlas using the function IntegrateLayers and the method “CCAIntegration”. Dimensional reduction was performed with the RunPCA function. The first 40 dimensions were then used to build a shared nearest-neighbor network with the function FindNeighbors and for visualization in UMAP space with the function RunUMAP. Cells were clustered using the Louvain algorithm from the function FindClusters. We clustered at two resolutions (0.2 and 0.8) resolutions to enable analysis of major cell types and their subpopulations^8^.

### Ortholog identification

We employed a reciprocal best hit approach to identify *Drosophila melanogaster* orthologs for *Polistes fuscatus* genes. Proteins for each species were queried against each other in BLASTP (blast 2.16.0) searches with E value thresholds of 1 x 10^-5^. Protein pairs that were the best hits in both directions were considered putative orthologs. The same approach was used to identify orthologs from *Harpegnathos saltator*, *Apis mellifera*, and *Monomorium pharaonis*. *Drosophila* (BDGP6.54) proteins were downloaded from Ensembl. *Harpegnathos* (HSal_v8.6), *Apis* (HAv3.1), and *Monomorium* (ASM1337386v2) datasets were obtained from the National Center for Biotechnology Information (NCBI).

### Cell annotation and transcriptional similarity

Cell type annotation was performed using a combination of cluster-specific marker genes and cross-species transcriptional similarity (below). Marker genes were identified with the FindAllMarkers function and Wilcoxon Rank Sum tests. Cell types were assigned by overlapping significant marker genes (adjusted p values < 0.05, logFC > 0.25, and pct.1 > 0.2) with established markers from the literature. Marker gene identification was performed twice for the central brain atlas (resolution = 0.2, 0.8) and once for the whole brain worker atlas (resolution = 0.8).

We used MetaNeighbor v3.13^51^ to assess the transcriptional similarity of clusters across species. MetaNeighbor estimates the similarity of cell clusters across datasets according to the expression of shared genes, enabling annotation of cell types and estimation of similarity across species. Published single-cell datasets from *D. melanogaster*^10^, *H. saltator*^14^, *M. pharaonis*^13^, and *A. mellifera*^16^ were analyzed against the *P. fuscatus* central and whole brain atlas. To increase reliability and reduce noise inherent in single-cell datasets, we examined transcriptional similarity with pseudo-cells, as done in previous studies^13,100^. Within each cluster, expression values of 10 randomly drawn cells were merged to create pseudo-cells. Expression was summed by gene, converted to counts per million, then standardized with z-transformation. Interspecies cluster similarity was then quantified using MetaNeighbor and represented with pairwise AUROC scores. Atlas-wide contrasts only included genes that were significant markers for clusters in at least one dataset. Kenyon cell contrasts had a limited number of shared marker genes, so we lessened our restrictions to include any orthologous genes shared across all datasets.

### Differential abundance and expression across phenotypes

Cell abundance in single-cell datasets is compositional in nature, meaning linear models on counts and proportions are insufficient for testing differential abundance of cell clusters. Here, we assessed variation in cell abundance via a Bayesian framework within scCODA v0.1.9^101^, which explicitly models compositional changes. We quantified divergence in primary and secondary cluster abundance across the life stages via the CompositionalAnalysis function with Hamiltonian Monte Carlo sampling. Because we had no a priori expectations about cell abundances, an automatic reference cell type selection was used. Differential abundance was examined across each pairwise stage contrast and significance was called at FDR < 0.1, a non-zero scCODA final parameter, and a log2 fold change greater than 0.38^13^. scCODA uses a Bayesian variable-selection framework to control false discovery rate, so no additional multiple-testing corrections were applied. These thresholds are conservative, as scCODA can shrink small effect sizes and under-detect subtle compositional shifts, particularly in clusters with small cell counts.

Patterns of differential expression were examined at two levels. First, differentially expressed genes were identified using the Seurat FindMarkers function. The number of biological replicates in each group was limited, meaning pseudobulk analyses were not feasible, so differential expression was instead quantified using cell-level tests within clusters. Stage-specific differential expression was calculated for each pairwise contrast using MAST models, which account for variability in transcript detection across cells. We considered genes biologically meaningful with an adjusted p value < 0.05 and log2 fold change > 0.5. Next, we used hierarchical clustering to estimate the transcriptional similarity of phenotypes within each cluster and across the entire central brain. For each cluster and library, we aggregated transcript counts to generate pseudobulk profiles and logCPM normalized the values. Low variance genes can introduce noise, so we retained the 3,000 most variable genes and calculated Pearson correlation distance between libraries (represented as 1-*r*). To convert to condition-level distances, we collapsed the distance matrix to the median pairwise distance between samples in a contrast. Hierarchical clustering for plotting was performed with the function hclust from the base R *stats* package.

### Social interaction assay

Foundresses and their nests were collected from the wild and isolated for a minimum of two days prior to any behavioral assays. Wasps were placed in 3D printed arenas under three conditions. Paired “fighters” were placed together in the same chamber, “watchers” were placed in an adjacent chamber separated by a clear acrylic barrier, and “solitary” wasps were placed alone in a chamber without visual, physical, or olfactory cues of other wasps. Wasps were sacrificed and dissected 45-60 minutes after the assay, following a previous study of socially mediated bulk gene expression in fighter wasp brains^37^.

Single-cell libraries were sequenced and integrated as described above. Cluster-wise differential expression was calculated using Wilcoxon Rank Sum tests in the Seurat FindMarkers function. Genes were considered differentially expressed relative to the “solitary” control condition at an adjusted p value < 0.1. DEG overlap among fighters and watchers was assessed using a hypergeometric enrichment test from the function phyper. Significance was defined according to FDR-corrected p-values < 0.01 and an overlap greater than three genes. Patterns of differential expression were similar for primary and secondary clusters, so we report results from secondary clusters for greater resolution.

### IEG detection and neural activation

In insects, four genes exhibit rapid transcription following neural activation (*sr*, *hr38*, *Jra*, *kayak)*^69^. For each of these canonical IEG, cells were designated as IEG-positive or negative. Cells were designated IEG-positive when their log-expression exceeded 1, a threshold selected to reduce background signal and ensure accurate detection of activated cells. We then calculated differential expression between positive and negative cells within each cluster to identify genes that differed between activated and non-activated cells. This approach identified genes that were significantly co-expressed with canonical IEGs, which could therefore be used as an expanded panel to measure neural activation. Differential expression was called via the FindMarkers function and required a minimum of 3 cells per group to call DE (as per minimum requirements for FindMarkers). Genes were considered significant with an adjusted p value < 0.05 and a gene was classified as IEG-like if it was significantly associated with a canonical IEG in the majority of neural clusters. Non-neural clusters are not expected to exhibit activation patterns and were therefore excluded from IEG panel discovery. These steps were performed separately for each canonical IEG. Both *hr38* and *sr* exhibited significant co-expression but were not co-expressed with *Jra* or *kayak,* in part because *kayak* exhibited limited expression in our dataset. We therefore required IEG-like genes to exhibit co-expression with both *hr38* and *sr* in our analyses. Though patterns of neural activation in response to social interactions were only tested in foundress conditions, we repeated the IEG panel discovery separately for each caste. Six genes were shared across all sexes and castes, all of which were identified in the foundress-only dataset, supporting the validity of the approach (Fig. S11b). We opted to use the larger foundress panel for downstream analyses due to their similar collection strategy and timing.

To examine signatures of activation across clusters, we used the panel of IEG-like genes to assign activation scores to individual cells. A similar approach in cichlids defined activation according to the number of IEGs expressed in a cell^8^. To account for variation in expression levels of each IEG, we instead used Seurat’s AddModuleScore function, which generates a score that incorporates both the expression of panel genes and background expression of binned random control genes. Because single-cell datasets contain substantial noise, we considered a cell “activated” if its activation score was in the top 20% of the distribution.

To identify activated subpopulations of cells, we calculated differential expression between activated and non-activated cells within each cluster^70^. Significant genes indicated cell populations that consistently distinguish activated and non-activated states, and therefore identified cluster subpopulations that activate in response to stimuli. Conversely, a lack of activation-related DEGs suggested that subpopulations of cells have no consistent response to stimuli. Our limited biological replicates precluded pseudobulk analyses, so differential expression was quantified using the FindMarkers function with the MAST test, which is designed for zero-inflated single cell data and incorporates detection rate and continuous expression in the model. We called differential expression within each condition at adjusted p value < 0.1 and examined secondary clusters to provide finer resolution of gene-defined subpopulations. To focus on socially-responsive genes, we excluded genes associated with activation in the solitary condition. DEG overlap between the fighter and watcher conditions, and among neural clusters, was assessed via hypergeometric enrichment tests and significance was defined according to FDR-corrected p-values < 0.01 and an overlap greater than three genes.

## Author Contributions

M.J.S. conceived and designed the study. C.M.J. and M.C. performed experiments, dissections, and sample preparation. C.B. analyzed and visualized data. C.B. and M.J.S. wrote the manuscript with input from M.C., C.M.J., R.D.R, and A.L.T.

## Data Availability

Raw single-cell RNAseq data from *P. fuscatus* generated in this study are deposited in the NCBI Sequence Read Archive under BioProject accession PRJNA1546199 (under controlled access until publication). Raw PacBio IsoSeq reads used for improving the *P. fuscatus* annotation are deposited under BioProject accession PRJNA1545971. Processed single-cell datasets, analysis objects, and supplementary data will be deposited in a public repository upon publication.

