## Supplemental Figures for "A single-cell transcriptomic atlas of paper wasp brains highlights Kenyon cells’ role in social behavior"

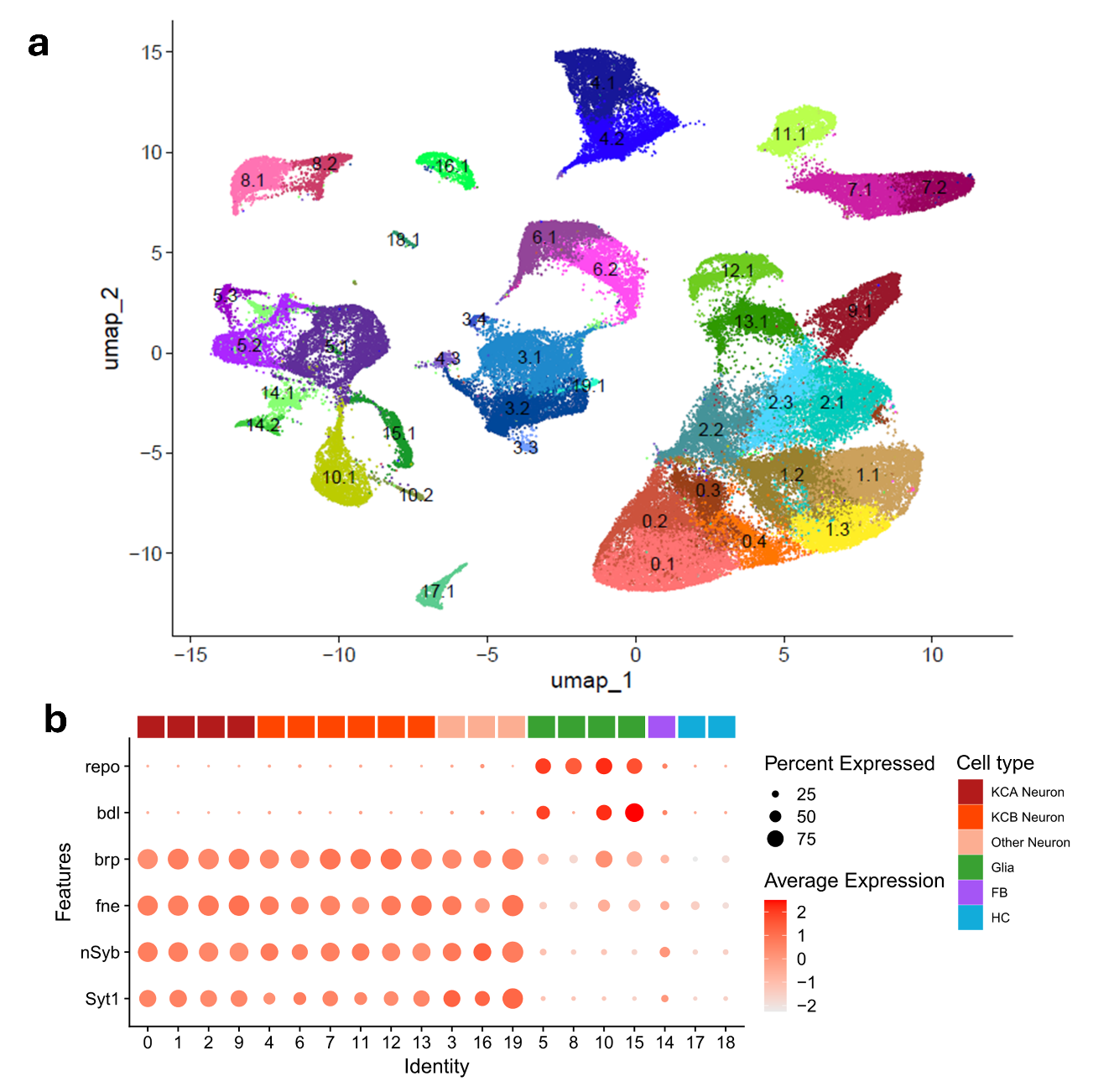

**Figure S1. Characterizing cell populations of the central brain. a.** UMAP plot of the 39 secondary clusters (resolution 0.8) comprising the cell atlas. Each point represents a single cell of the 96,310 cell atlas. Subclusters are named according to their assignment from primary clustering. **b.** Neural and glial cells are cleanly separated according to marker gene expression. The dot plot displays the expression of major glial (*repo, bdl*) and neural (*brp, fne, nSyb, Syt1*) marker genes across cell clusters. Broad cell types are annotated with colored bars.

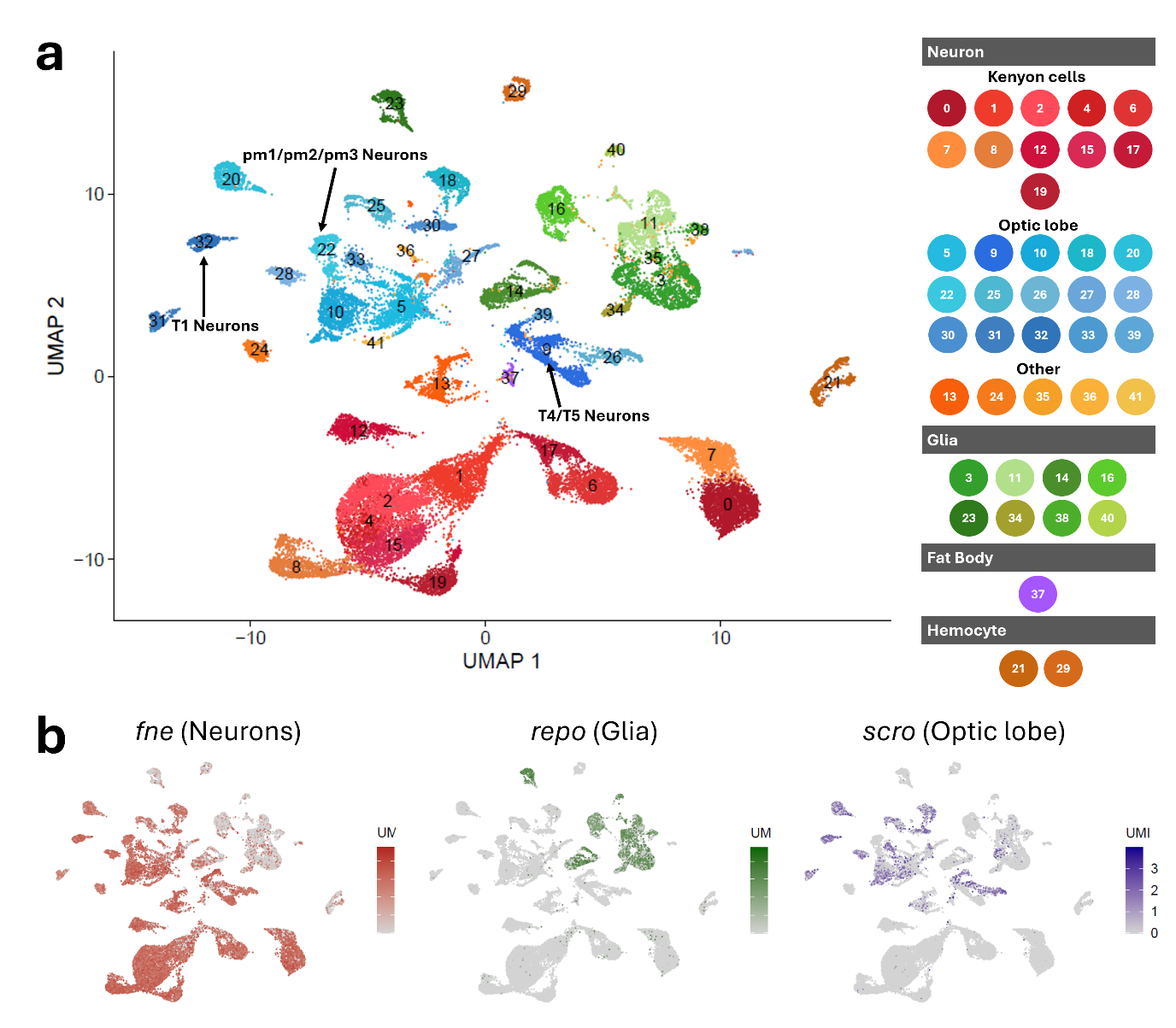

**Figure S2: The whole-brain atlas of a *Polistes fuscatus* worker.** **a.** UMAP plot of the 42 clusters comprising the combined optic lobe and central brain cell atlas. Each point represents a single cell of the 33,439 cell worker atlas. The legend indicates major groupings and known cell types of clusters, with specific optic lobe clusters labelled with arrows. **b.** Marker gene expression of whole brains visualized in UMAP space. The gene *fne* is a marker for neural cells, *repo* is a marker for glial cells, and *scro* is a marker for many optic lobe neural cells.

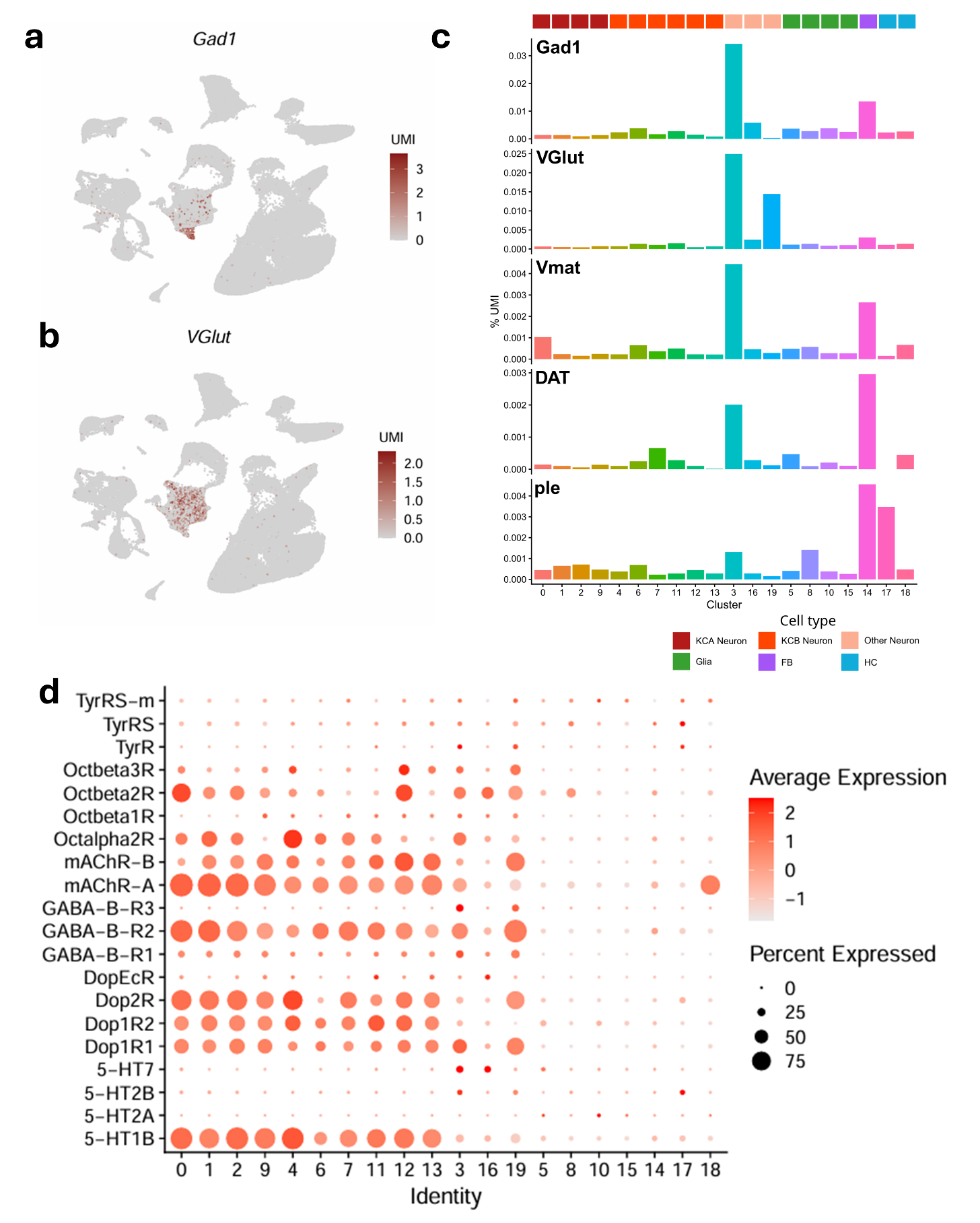

**Figure S3. Neuropeptide expression in the central brain. a-b.** Expression of GABAergic (*Gad1*) and glutamatergic (*VGlut*) marker genes is largely restricted to cluster 3 neurons. Expression levels are visualized in UMAP space. **c.** Neurotransmitter pathway genes are primarily expressed in neural cluster 3 and fat body cluster 14, suggesting that the majority of neurotransmitter synthesis occurs in these clusters. **d.** Neurotransmitter receptors are broadly expressed across neural clusters, particularly Kenyon cells. The dot plot displays average expression and percent-expressing cells for each receptor across central brain clusters.

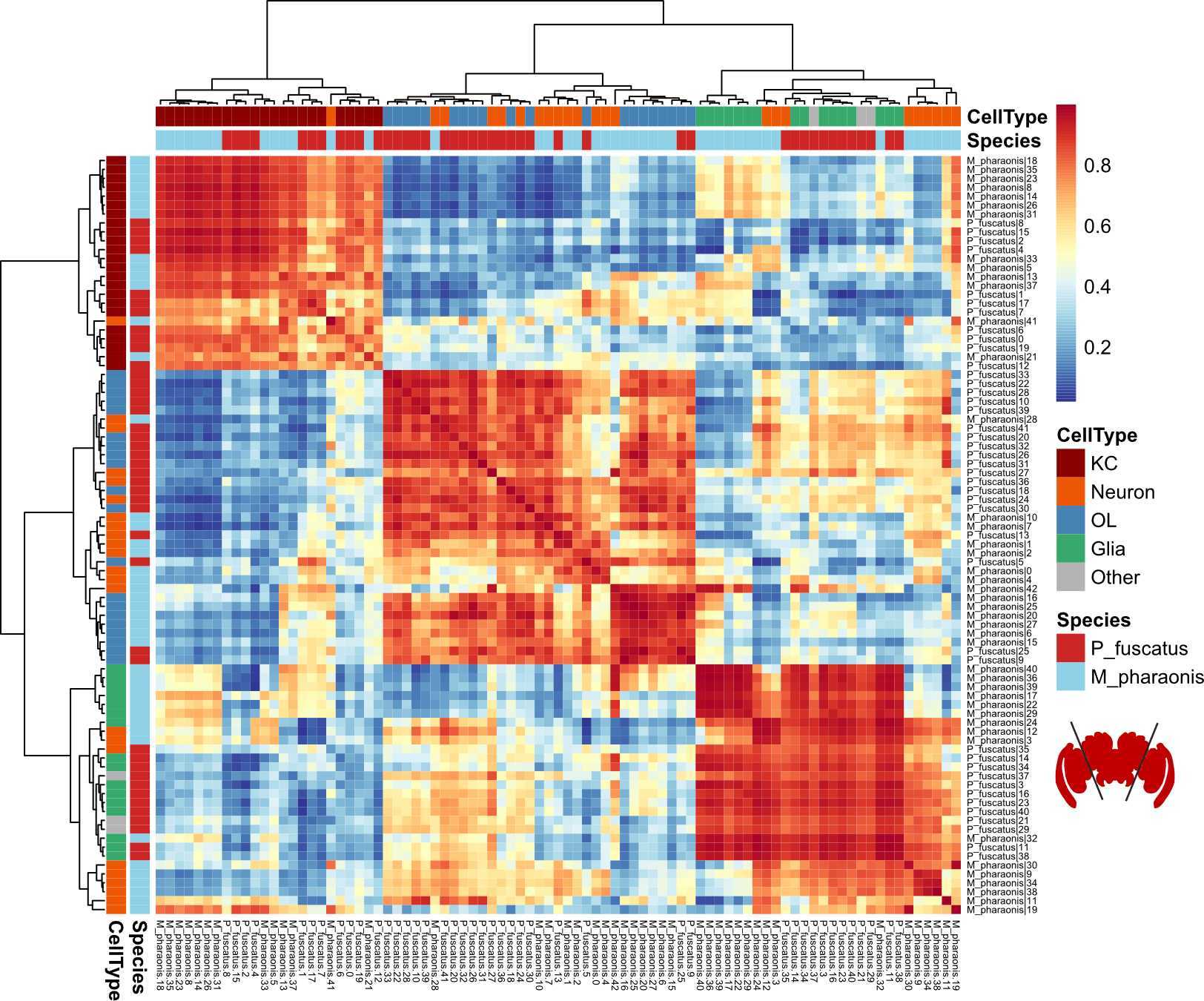

**Figure S4. *Polistes* brain atlas classification is robust against the *Monomorium pharaonis*** **brain atlas.** Pairwise transcriptional similarity of whole brain cell clusters from *Polistes fuscatus* against the pharaoh ant *Monomorium pharaonis* from MetaNeighbor AUROC scores. Kenyon cells, glia, and optic lobe neurons consistently clustered together across species. Columns and rows are annotated according to the species and cell type. KC, Kenyon cell; OL, Optic lobe neuron.

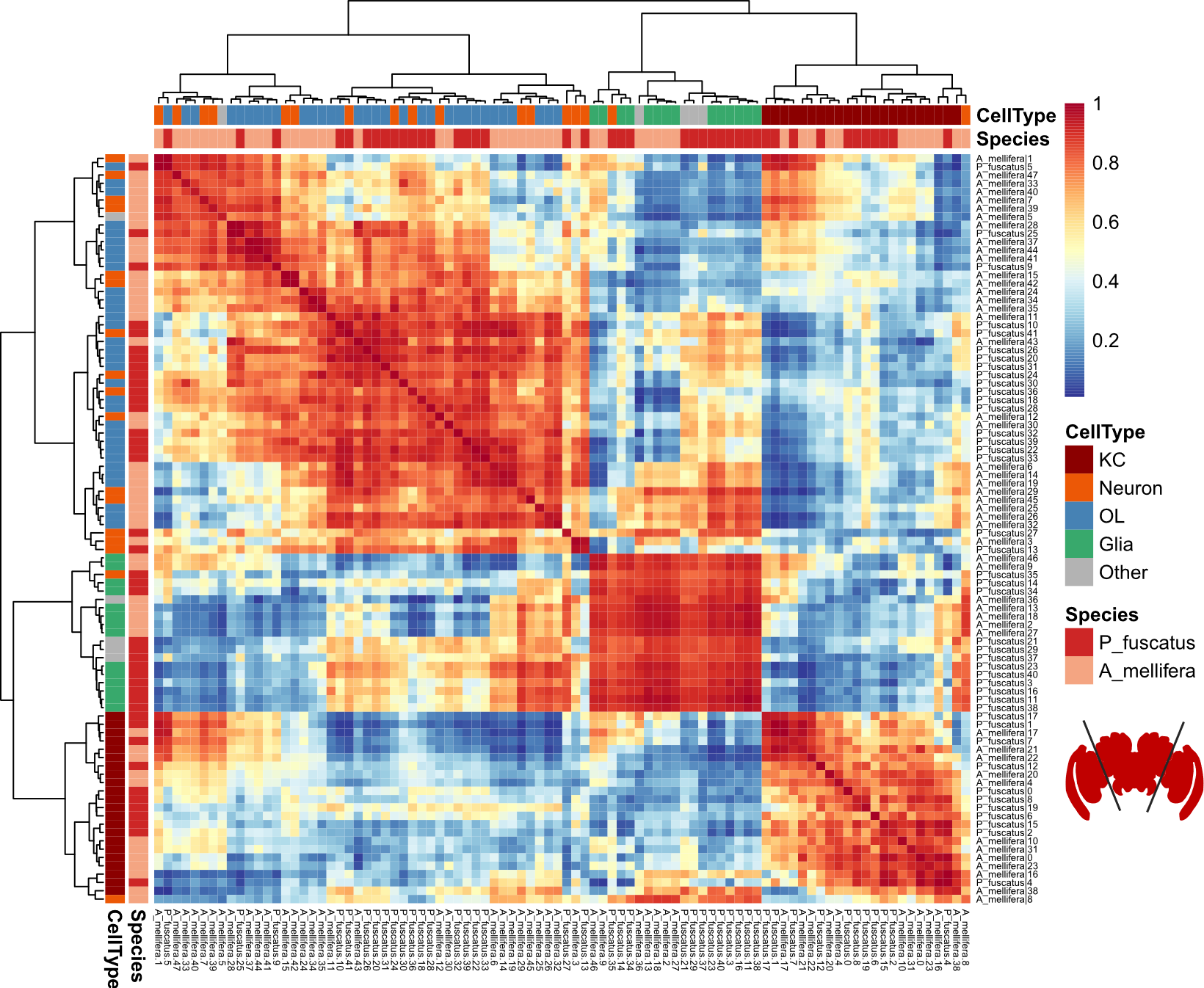

**Figure S5. *Polistes* brain atlas classification is robust against the *Apis mellifera* brain atlas.** Pairwise transcriptional similarity of whole brain cell clusters from *Polistes fuscatus* against the honey bee *Apis mellifera* from MetaNeighbor AUROC scores. Kenyon cells, glia, and optic lobe neurons consistently clustered together across species. Columns and rows are annotated according to the species and cell type. KC, Kenyon cell; OL, Optic lobe neuron.

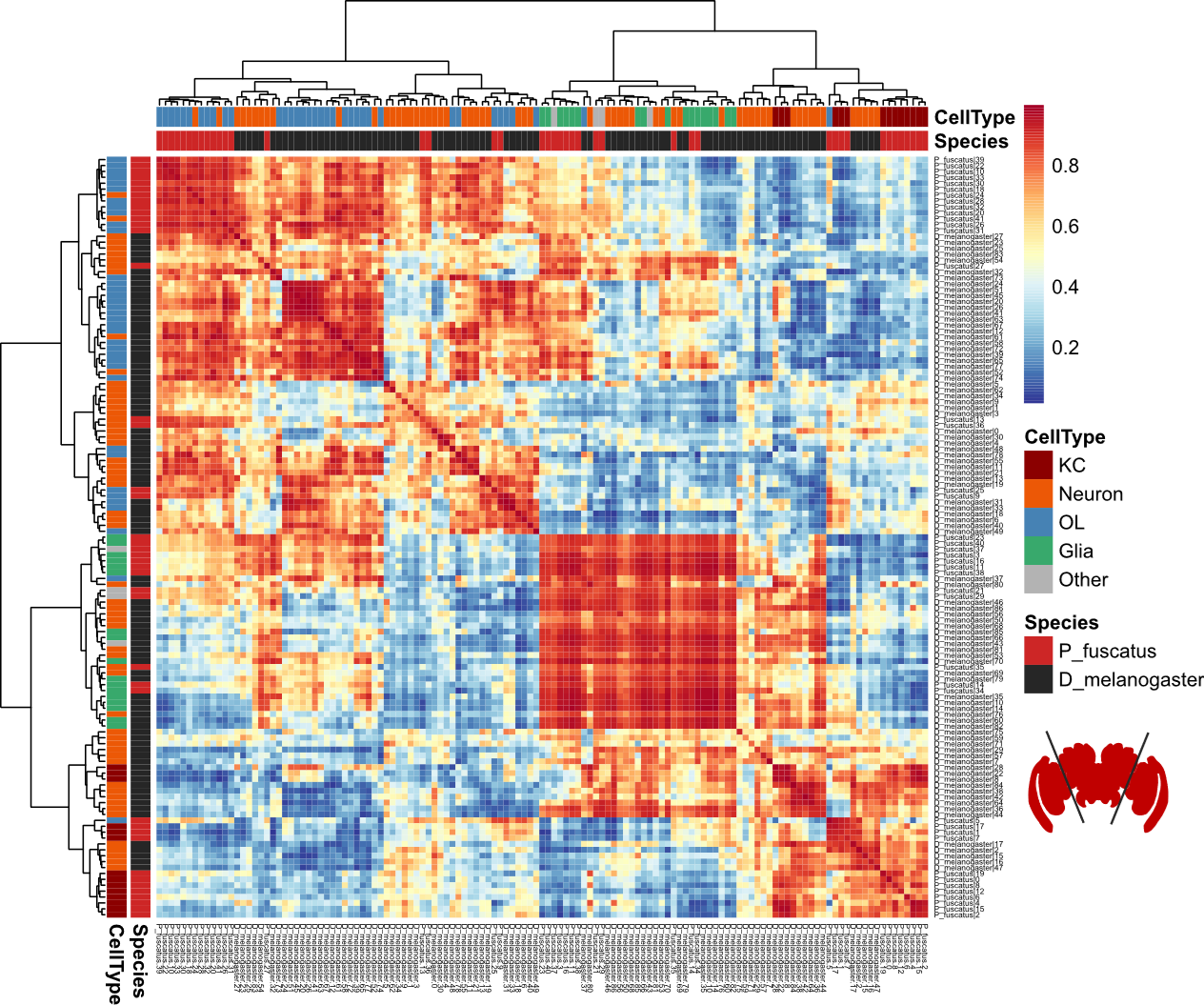

**Figure S6. *Polistes* brain atlas classification is robust against the *Drosophila melanogaster* brain atlas.** Pairwise transcriptional similarity of whole brain cell clusters from *Polistes fuscatus* against *Drosophila melanogaster* from MetaNeighbor AUROC scores. Kenyon cells, glia, and optic lobe neurons consistently clustered together across species. Cluster relationships were notably weaker than the Hymenoptera-specific contrasts, which is expected due to their evolutionary distance. Columns and rows are annotated according to the species and cell type. KC, Kenyon cell; OL, Optic lobe neuron.

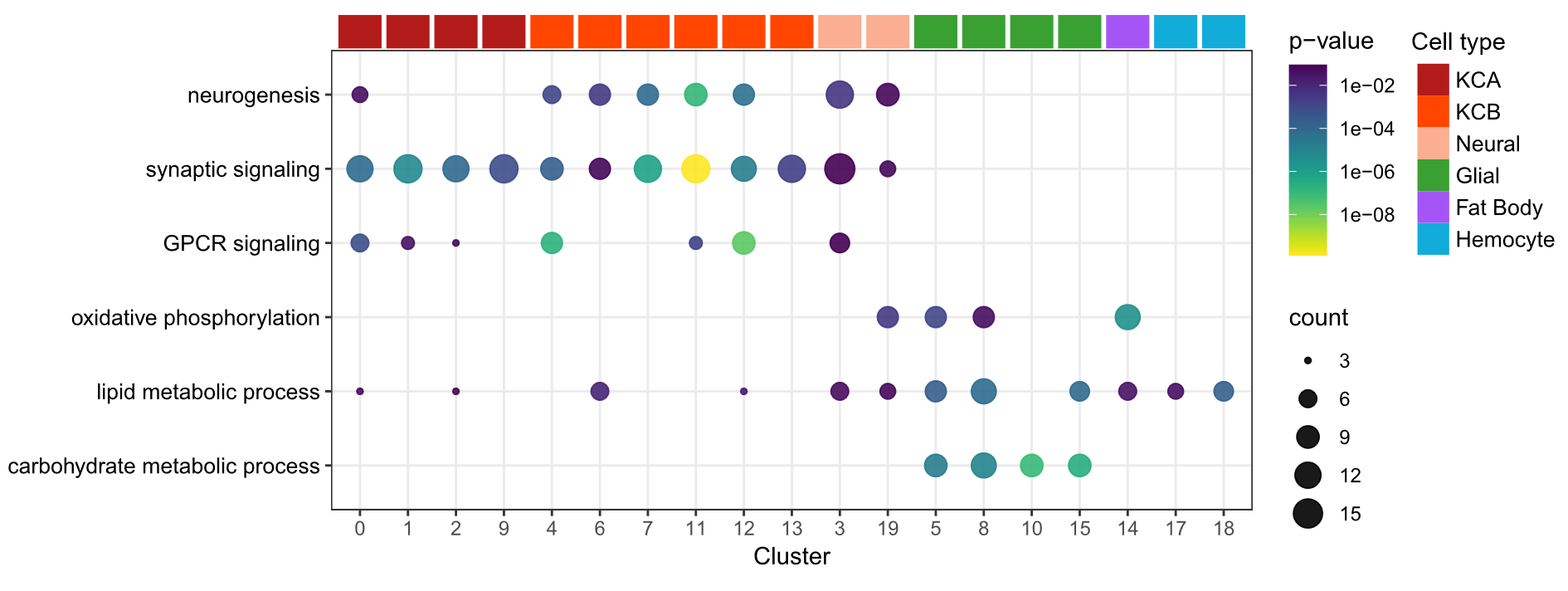

**Figure S7. Representative GO terms reflect cluster identities.** In support of cluster annotations, neural cluster markers are enriched for neural signaling processes while glial cluster markers are enriched for metabolic processes. Significantly enriched (p < 0.1) markers are colored by p value and sized according to the number of DEGs associated with each GO term.

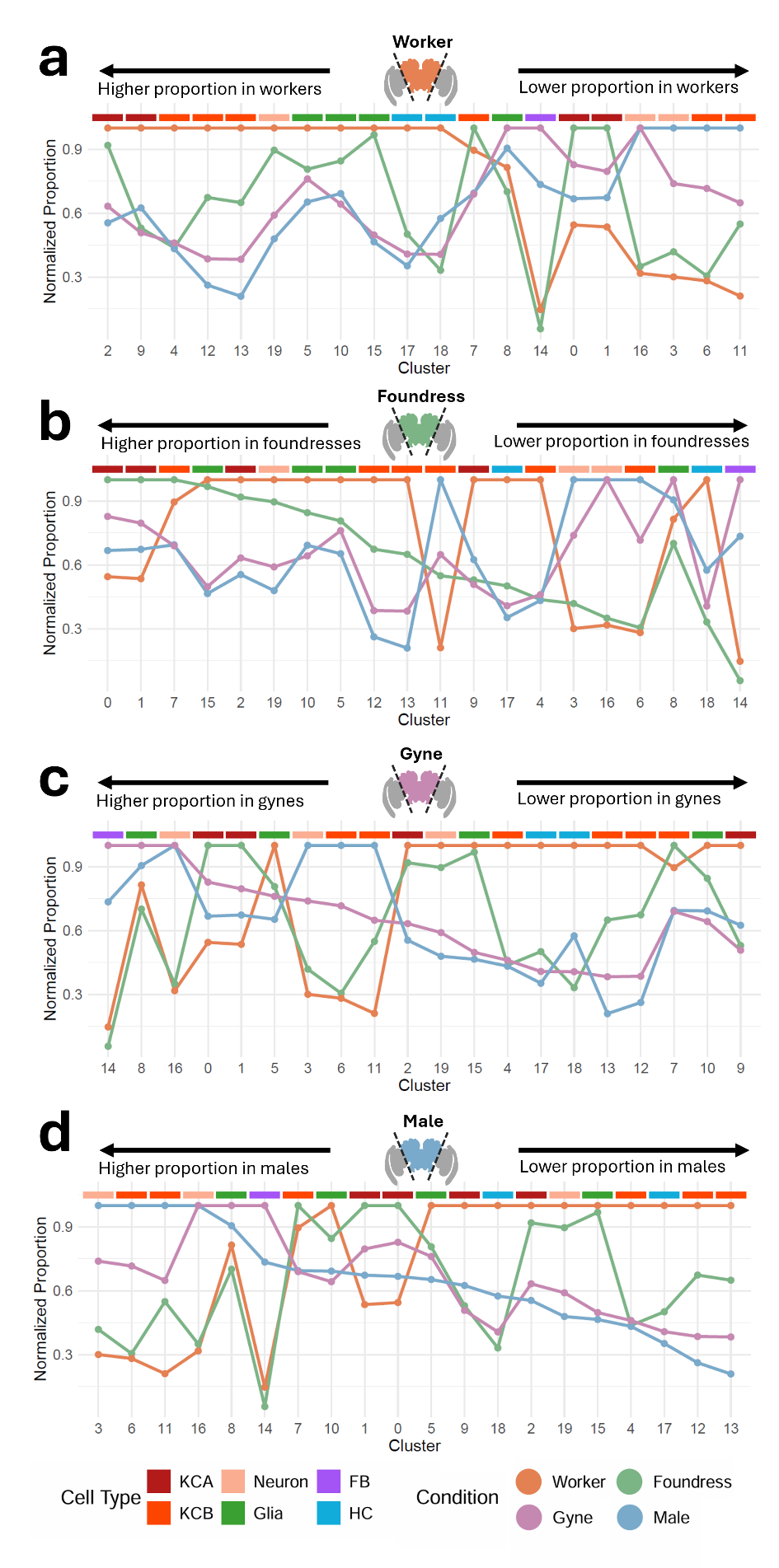

**Figure S8. Specialization of wasp brain composition across phenotypes.** The relative abundance of each cell cluster is plotted across phenotypes. For each cluster, the proportion of cells has been normalized across phenotypes to the maximum value among groups. Clusters are ordered according to abundance in workers **(a)**, foundresses **(b)**, gynes **(c)**, and males **(d)**. Worker cell proportions were often the most extreme compared to other castes, suggesting worker brain compositions are highly specialized, while cell proportions in gynes and foundresses were often moderate.

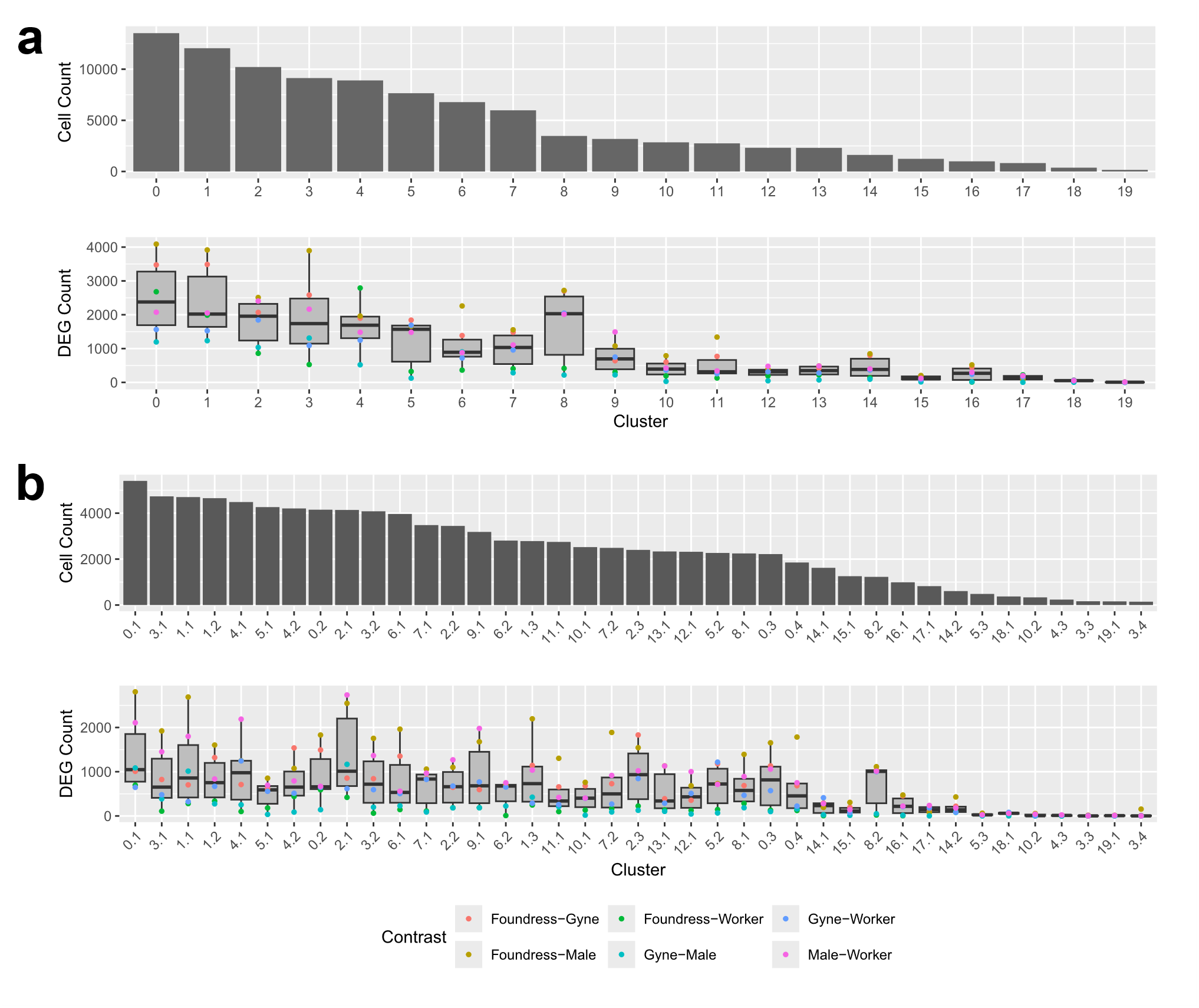

**Figure S9. Detection of differential expression scales with cluster size. a.** Primary cell clusters vary significantly in size. Power to detect differential expression is closely-associated with cluster size, which confounds comparisons of transcriptional dynamics across clusters. **b.** Secondary clusters vary less in size, enabling comparisons of differential expression across similarly-sized clusters and cell types.

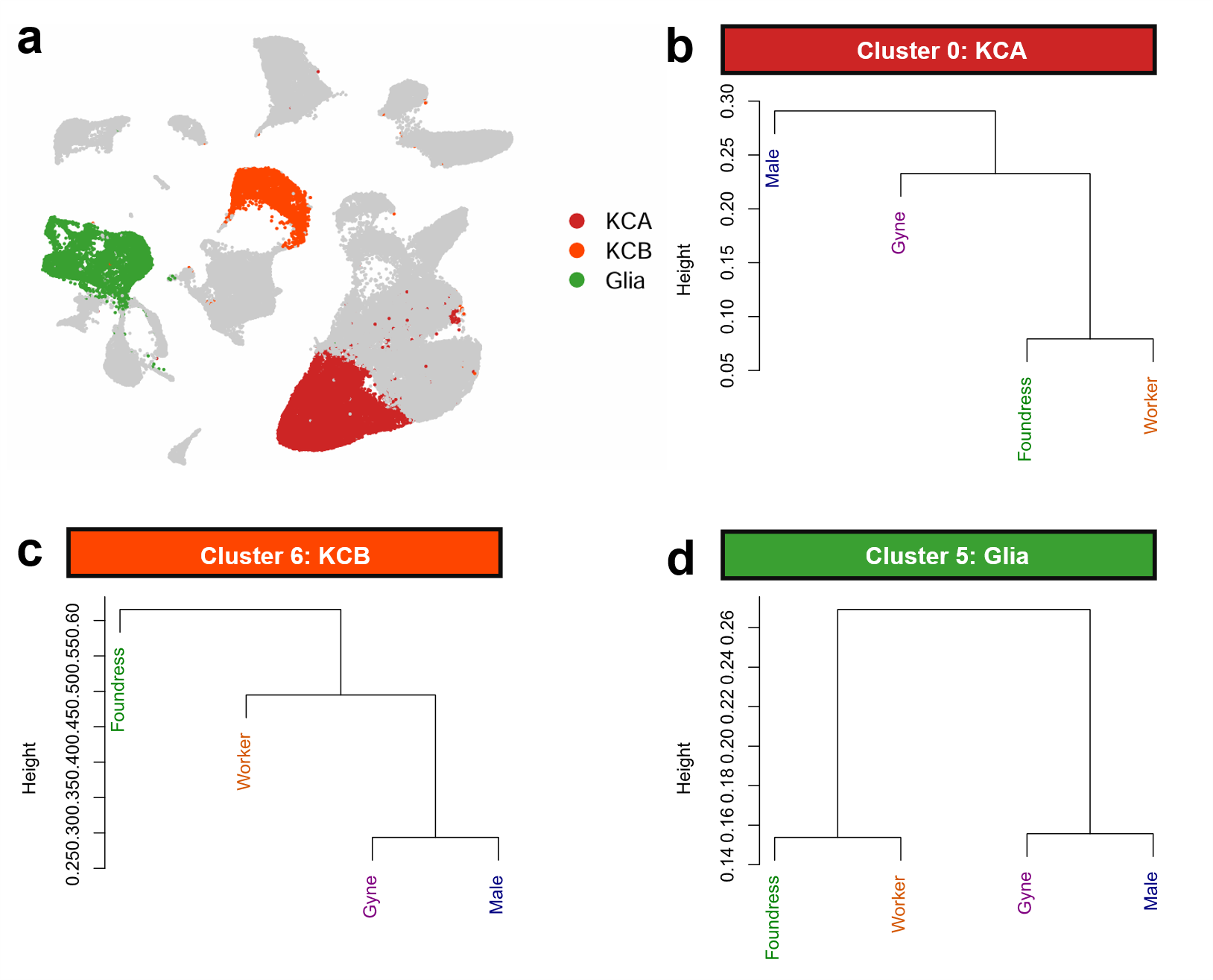

**Figure S10. Clustering relationships among life stages is heterogenous across cell types.** Although pseudobulk analysis suggests that transcriptomic profiles cluster primarily according to season (Fig 4d), cluster-specific patterns are hetergeneous. **a.** Example clusters are highlighted in UMAP space. **b.** Class A Kenyon cell cluster 0 groups transcriptomic profiles according to sex, with males in the outgroup. **c.** Class B Kenyon cell cluster 6 places foundresses in the outgroup, suggesting that expression patterns in this cluster may be associated with the behavioral and reproductive demands of early nest founding. **d.** Glial cluster 5 separates phenotypes according to season. Early-season foundresses and workers cluster separately from late-season gynes and males, potentially reflecting shifts in metabolic demands throughout the breeding season.

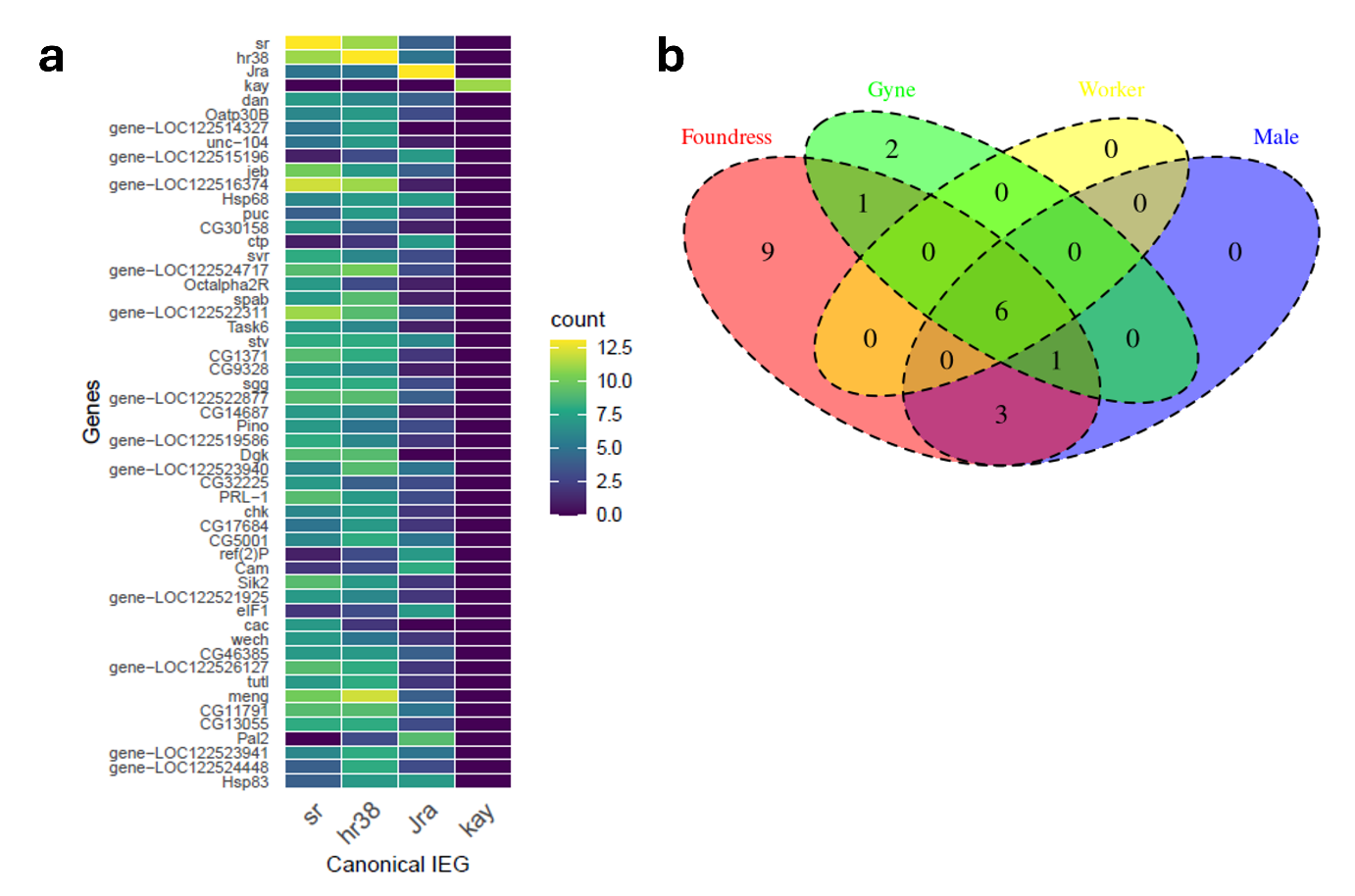

**Figure S11. Immediate early genes nominate a panel of co-expressed genes. a.** Genes co-expressed with canonical IEGs from the foundress dataset. Genes coexpressed with *sr* and *hr38* are generally consistent, suggesting both canonical genes can be used to profile neural activation. *Jra* exhibited distinct expression patterns, while *kayak* was poorly expressed in this dataset. Tiles are colored according to the number of neural clusters with significant co-expression. **b.** Venn diagram of IEG-like genes (as defined by co-expression with *hr38* and *sr*) nominated across each life stage dataset. Six IEG-like genes were shared across life stages, indicating that this approach captures consistent signatures of activation.
